# Single-nuclei RNA-sequencing of plant tissues

**DOI:** 10.1101/2020.11.14.382812

**Authors:** Daniele Y. Sunaga-Franze, Jose M. Muino, Caroline Braeuning, Xiaocai Xu, Minglei Zong, Cezary Smaczniak, Wenhao Yan, Cornelius Fischer, Ramon Vidal, Magdalena Kliem, Kerstin Kaufmann, Sascha Sauer

**Author notes:** Joint Authors (Daniele Y. Sunaga-Franze, Jose M. Muino, Caroline Braeuning, Xiaocai Xu).

## Abstract

Single-cell genomics provides unprecedented potential for research on plant development and environmental responses. Here, we introduce a generic procedure for plant nuclei isolation combined with nanowell-based library preparation. Our method enables the transcriptome analysis of thousands of individual plant nuclei. It serves as alternative to the use of protoplast isolation, which is currently a standard methodology for plant single-cell genomics, although it can be challenging for some plant tissues. We show the applicability of our nuclei isolation method by using different plant materials from several species. The potential of our snRNA-seq method is shown through the characterization of transcriptomes of seedlings and developing flowers from *Arabidopsis thaliana*. We evaluated the transcriptome dynamics during the early stages of anther development, identify stage-specific transcription factors regulating this process and the prediction of their target genes. Our nuclei isolation procedure can be applied in different plant species and tissues, thus expanding the toolkit for plant single-cell genomics experiments.

**SIGNIFICANCE STATEMENT:** We introduce an optimized plant nuclei isolation procedure followed by single nuclei RNA-seq that can be applied to different plant tissues without the need for protoplast isolation.

## INTRODUCTION

The fundamental units of life, the cells, can vary tremendously within an organism. The analysis of specialized cells and their interactions is essential for a comprehensive understanding of the function of tissues and biological systems in general. Major biological roles such as growth, development and physiology ultimately gain plasticity from heterogeneity in cellular gene expression (Fischer *et al*., 2019).

Without precise transcriptional maps of different cell populations, we cannot accurately describe all their functions and underlying molecular networks that drive their activities. Recent advances in single-cell (sc) and in particular single-nucleus (sn) RNA-sequencing have put comprehensive, high-resolution reference transcriptome maps of mammalian cells and tissues on the agenda of international consortia such as the Human Cell Atlas (Regev *et al*., 2017).

Similar efforts are made by the Plant Cell Atlas (Rhee *et al*., 2019). Plant tissues and plant cells pose specific challenges compared to mammalian systems (Efroni and Birnbaum, 2016). Plant cells are immobilized in a rigid cell wall matrix, which is required to be removed for isolating single cells. Additional technical demands include size variability of plant cells and the presence of plastids and vacuoles. Consequently, these characteristics require considerably different operational procedures compared with mammalian tissues. Recently, plant single-cell RNA-sequencing studies using protoplast isolation (PI) have been published (e.g. Efroni *et al*., 2015; Efroni *et al*., 2016; Nelms and Walbot, 2018; Zhang *et al*., 2019; Jean-Baptiste *et al*., 2019; Denyer *et al*., 2019; Shulse *et al*., 2019; Ryu *et al*., 2019). This procedure allows to sensitively identify and classify plant cell types. However, it is known that enzymatic digestion of plant cell walls can introduce artifacts at the transcriptome level, limiting the applicability of this approach (Shulse *et al*., 2019; Jean-Baptiste *et al*., 2019). To overcome this limitation, PI-response genes can be identified through an independent bulk RNA-seq experiment and later eliminated from the scRNA-seq analysis (Denyer *et al*., 2019).

Several methodologies are available for the generation of RNA-seq libraries from isolated cells or nuclei. Two of the most popular are: droplet-based (e.g. 10x Chromium) and nanowell-based (e.g. Takara iCELL8) systems. Droplet-based methods are popular because of their scalability. They enable rapid processing of thousands of cells simultaneously. Particularly in the Chromium system, gel beads are supplied with a unique barcoded oligonucleotide. Cells or nuclei are encapsulated together with these beads, lysed and the gel bead releases the barcoded oligonucleotide for reverse transcription (RT). RT is performed inside droplets and transferred to a tube where amplification of cDNA occurs. One disadvantage is that, in some events, more than one cell or nucleus enters the same capsule, producing a mixed cDNA population (Lareau *et al*., 2020). Nanowell-based systems trap the isolated cells or nuclei in wells where the cDNA is produced. In particular, iCELL8 system consists of a chip with more than 5,000 nanowells containing barcoded oligonucleotides attached to their surface. Each cell or nucleus is deposited in a nanowell and its quality and number are checked by microscope, which reduces the probability of obtaining transcriptomes from more than one cell/nucleus. One of the main disadvantages of nanowell systems is their more limited scalability, as each chip has a fixed number of nanowells. To our knowledge, only the droplet-based systems have been applied in plants, however it is crucial to continue enriching and improving the repertoire of single cell omics methodologies available for the plant research community.

Here, we introduce a single-nucleus sequencing protocol using the nanowell-based iCELL8 system by studying the dynamics of *Arabidopsis* transcriptomes during flower development. Working with nuclei has the advantage of eliminating organelles and vacuoles, as well as secondary metabolites localized in the cytoplasm that can interact with RNA and negatively affect NGS library preparation.

## RESULTS AND DISCUSSION

### Nuclei isolation and snRNA-seq library preparation

While protoplast isolation (PI)-based methods have been shown to readily identify plant cell types, it is also known that it can lead to changes in gene expression and different cells types may be affected in different degrees (Shulse *et al*., 2019; Jean-Baptiste *et al*., 2019). To address this issue, PI responsive-genes can be identified through an independent bulk RNA-seq experiment and subsequently eliminating them from the scRNA-seq analysis (Denyer *et al*., 2019). However, we show that PI impact cannot be completely eliminated in this way (Supplementary Fig. 1a,b).

Here, we propose a single-nucleus sequencing (snRNA-seq) strategy for the transcriptome sequencing of individual plant nucleus (Fig. 1a; full protocol in Materials and Methods). The key step of our plant-nuclei sequencing procedure consists of gentle but efficient isolation of plant nuclei. Snap-frozen *Arabidopsis* tissue was gently physically dissociated by pestle and transferred to Honda buffer for cell lysis (Moreno-Romero *et al*., 2017). Cell walls and cell membranes were mechanically disrupted using a gentleMACS Dissociator, keeping the nuclei largely intact as observed by DAPI staining (Supplementary Fig. 2a). Released intact *Arabidopsis* nuclei were collected using Fluorescence-Activated Cell Sorting (FACS; Supplementary Fig. 2b). A clear separation between nuclei and debris was obtained (Supplementary Fig. 2c). To show the applicability of this method to different plant species/tissues, we successfully performed nuclei isolation in *Arabidopsis thaliana* (seedlings and flowers), *Petunia hybrida* (flowers), *Antirrhinum majus* (flowers), and *Solanum lycopersicum* (flowers and leaves) (Supplementary Fig. 2a,b). The RNA that was isolated from these nuclei was of high quality as observed by electrophoresis for *Arabidopsis* (Supplementary Fig. 2d).

**Figure 1:**
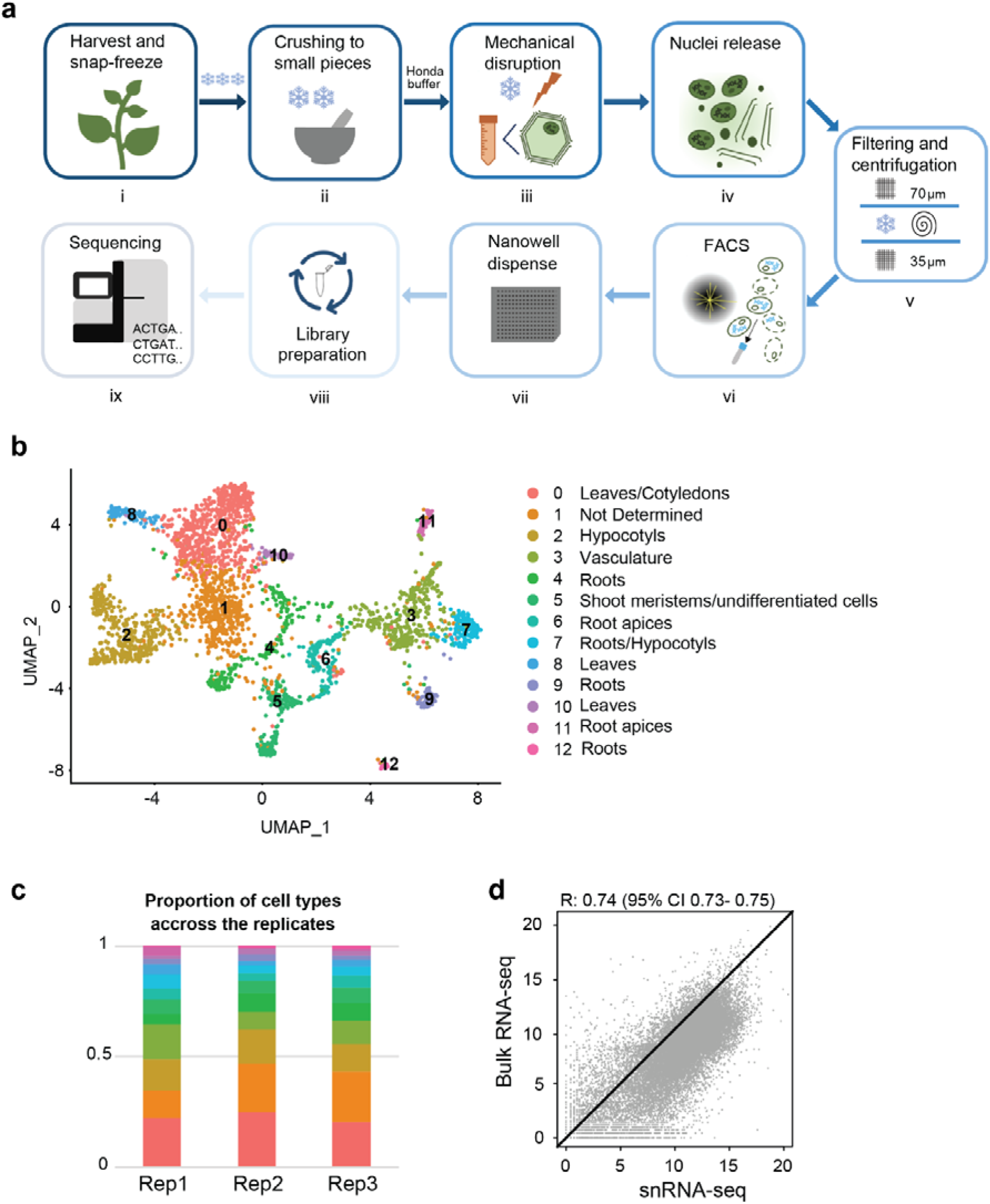
Single-nuclei RNA-sequencing. a) Schematic overview of snRNA-seq experimental strategy b) UMAP plot and clustering analysis of Arabidopsis seedlings samples (3 biological replicates, 13 clusters, 2,871 nuclei in total). c) Barplot showing that the three replicates have a similar proportion of nuclei across the identified clusters (the color code used to identify cluster cells is the same in b and c). d) Correlation (R= 0.74) of gene expression estimated from snRNA-seq (3 biological replicates) and bulk RNA-seq (3 biological replicates), indicating that snRNA-seq is able to recover similar transcriptomes than bulk RNA-seq.

The next step consists of generating high-quality cDNA libraries from the isolated nuclei. There is a number of different library preparation protocols and sequencing procedures that can be combined (9,22). We opted for the Takara’s ICELL8 system, a sensitive nanowell-based approach that includes a standardized lysis of nuclei by detergents and a freeze-thaw-cycle (Goldstein *et al*., 2017). One of the main advantages of this system is that it allows for manual selection of single-nucleus-containing wells, as well as visual inspection and selection of intact nuclei (i.e. nuclear rupture), thereby introducing additional quality control. Using SMARTer ICELL8 3’ chemistry, we prepared DNA libraries for short paired-end sequencing using fresh, snap-frozen *Arabidopsis* seedlings.

### snRNA-seq performance in Arabidopsis seedlings

To establish the method, we used 10-days-old *Arabidopsis thaliana* seedlings. Seedlings feature diverse plant structures, including the primary root, hypocotyl and cotyledons. This allowed us to characterize the performance of the method recovering the transcriptomes of diverse tissue types. A total of 3,348 nuclei was obtained from 3 biological replicates, with an average of 2,802 expressed genes per nucleus and 23,874 genes expressed in at least one nucleus per replicate (Supplementary Data 1, Supplementary Fig. 3). A Pearson correlation coefficient of 0.9 was observed among the biological replicates, indicating the high reproducibility of the method (Supplementary Fig. 4a,b). A good reproducibility was also observed between snRNA-seq and bulk RNA-seq (Pearson correlation of 0.74, Fig. 1d), even though snRNA-seq data comprise the nuclear transcriptome while bulk RNA-seq data comprise the nuclear and cytoplasmic transcriptome. It indicates that the method was able to recover the main transcript abundance present in the bulk RNA-seq data.

The integration of the 3 seedling datasets by Seurat revealed 13 major clusters (Fig. 1b). A similar proportion of nuclei from each annotated cluster was observed across the 3 replicates, again, indicating the good reproducibility of the method (Fig. 1c). To annotate the major tissue types enriched in each cluster, we first obtained the top 20 marker genes of each cluster (Supplementary Table 2). Then, the expression of these marker genes was characterized using a set of plant organ-specific bulk RNA-seq datasets (Transcriptome Variation Analysis Database; TraVaDB) (Supplementary Fig. 5). For example, cluster 12 has the highest signal in TraVaDB root samples, and therefore is labelled as “roots”. Since seedlings comprise a large diversity of cell and tissue-types, for many of which no cell-type specific transcriptome data are available, we did not pursue a comprehensive annotation of this dataset.

Since most published plant scRNA-seq experiments using PI-based methods focused on roots, we investigated the ability of our method to recover the main root cell types. We performed the re-analysis of a subset of 964 nuclei that were identified as “root” in our seedling snRNA-seq datasets (Fig. 1b, Supplementary Fig. 5). Twelve clusters were identified (Supplementary Fig. 6a). We calculated the overlap between the top 20 marker genes of each cluster (Supplementary Table 2) and the top 500 markers identified in a previous scRNA-seq study on root protoplasts (Denyer *et al*., 2019), revealing a clear overlap (Supplementary Fig. 6b). Sixty-five percent of the markers from cluster 11, for example, overlapped with the published cluster “10-xylem”. No overlap of these markers was found with any other cluster, indicating cluster 11 as a group of xylem-associated cells. Using this approach, we were able to recover all root cell types identified by (Denyer *et al*., 2019) except meristem, which might be explained by the low number of meristem cells in our “root” subset. The number of markers identified per cluster was lower in our case (ranging from 21 to 638) compared to Denyer *et al*. (ranging from 511 to 1,397). Possible reasons are the smaller number of root nuclei (964) compared to the comprehensive root atlas from Denyer *et al*., 2019 (4,727 cells) and the fact that our root nuclei were computationally selected from a pool of seedling nuclei. Despite these limitations, the results suggest that snRNA-seq can be used to identify cell types in complex samples, given the availability of cell-type-specific marker genes for annotation.

### Similarity between snRNA-seq data generated from fixed and unfixed plant material

To allow for more technical flexibility in our method, i.e. the possibility to simplify the storage of plant samples and maintaining *in situ* expression states (Alles *et al*., 2017), we fixed seedlings using methanol directly after harvest and performed snRNA-seq as described before. We obtained a similar number of nuclei (850) and an average number of expressed genes (2,292) when using methanol fixation compared to no fixation (1,116 nuclei and 2,802 genes). A similar nuclei distribution was also observed between fixed and non-fixed samples (Supplementary Fig. 7a). Additionally, an expression correlation of 0.88 and p-value<2.2e-16 was observed among both groups of samples (Supplementary Fig. 7b,c), indicating that fixation of the material does not introduce major differences in the number of nuclei and obtained cell-types.

### snRNA-seq performance in *Arabidopsis* inflorescences

To evaluate the performance of snRNA-seq to study cell differentiation, we applied snRNA-seq to *Arabidopsis thaliana* inflorescences, which cover all stages of flower development prior to anthesis. After quality control filtering, we obtained transcriptomes of 856 nuclei with an average number of 2,967 expressed genes per nucleus (Supplementary Fig. 8a). The analysis identified 15 clusters corresponding to distinct organs and developmental stages (Fig. 2a; Supplementary Fig. 8b). To annotate these clusters with particular cell types, we first identified specific marker genes of each cluster (Supplementary Table 2), then plotted their expression profiles in the different floral organs and developmental stages obtained from TraVaDB (Fig. 2b). Last, we correlated the gene expression of each cluster with each TraVaDB sample and indicated these values in the UMAP plot (Supplementary Fig. 8c). A major proportion of clusters (37% of the nuclei population) were annotated as differentiating anthers at different developmental stages (clusters 3, 4, 6, 7, 10, 15). This can be explained by the fact that anthers comprise a large fraction of tissues (Smyth *et al*., 1990; Gómez *et al*., 2015) in developing flowers. Furthermore, anthers/pollen have very specific gene expression profiles (Smyth *et al*., 1990; Gómez *et al*., 2015) which may facilitate the computational identification of the clusters. Our data captured gene expression dynamics during anther/pollen development from undifferentiated stem cells (cluster 0; Fig. 2) to late anther stages close to organ maturity, prior to anthesis (cluster 3; Fig. 2). This led us to use Monocle 3 to estimate the pseudotime of each anther cell (Supplementary Fig. 9c). When we plotted the average pseudotime of the cells of each anther cluster against the developmental time of each cluster obtained with the TraVaDB annotation (Supplementary Fig. 9d), it showed a strong concordance with anther developmental stages, which indicates that we can use the estimated pseudotime of each cell as a proxy of its developmental stage, and therefore to study transcriptional dynamics of anther differentiation.

**Figure 2:**
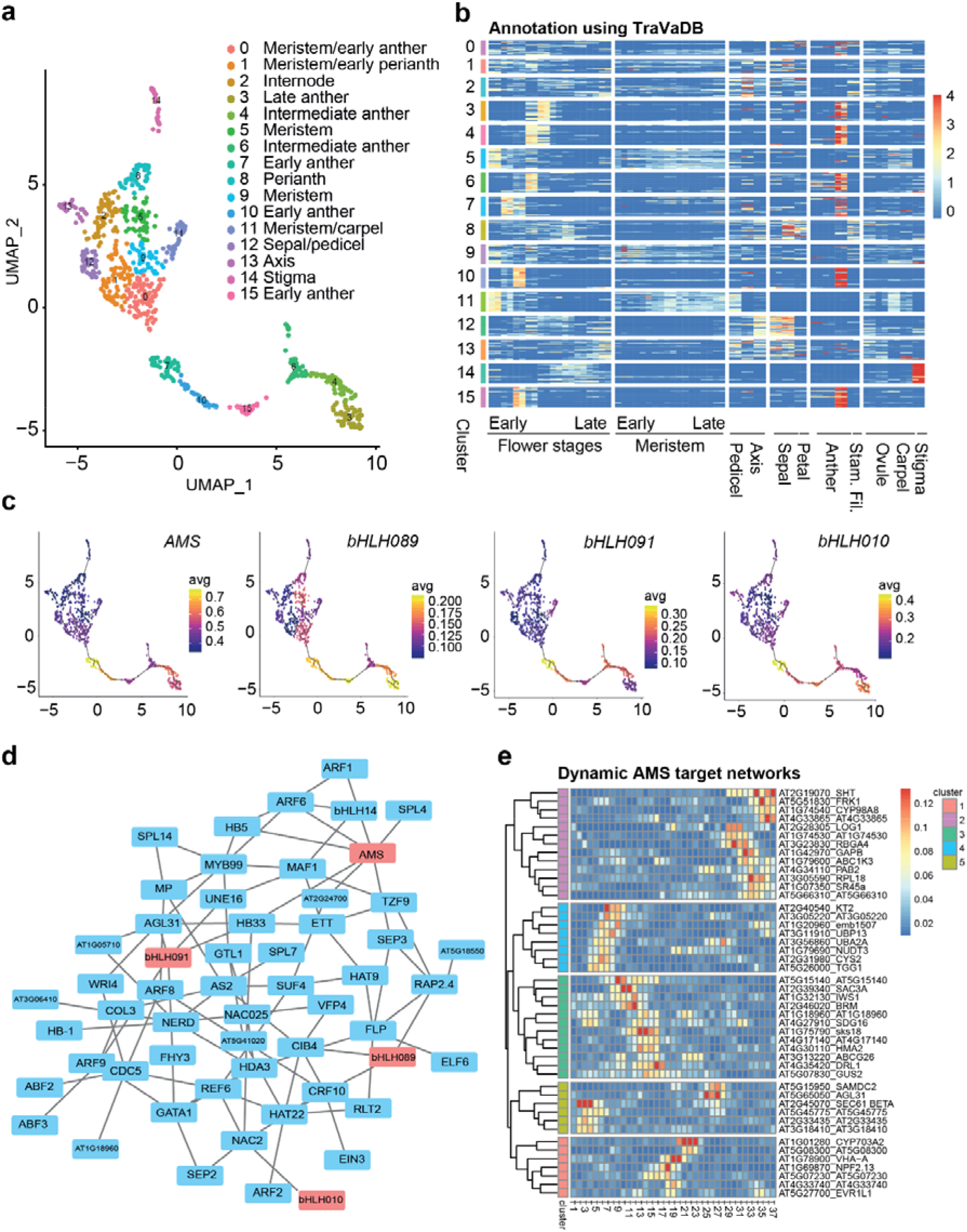
Anther development at single-nucleus resolution. a) UMAP plot and clustering of the snRNA-seq data from Arabidopsis flowers before anthesis. b) Heatmap showing the expression of the top 20 significant marker genes for each cluster. c) Gene expression of known representative anther TF regulators *AMS, bHLH089, bHLH0901* and *bHLH010* plotted in the UMAP coordinates. d) Gene network estimated from cluster 15 (early anther) using GENIE3 (only TFs with more than 3 targets are shown). e) Heatmap showing the strength of the interaction between AMS and its targets obtained by GENIE3 at different developmental stages. Cells belonging to an anther cluster were ordered by their developmental stage predicted by Monocle3 pseudotime analysis (Supplementary Fig. 9c) and GRN networks were predicted independently for overlapping sets of 50 cells ordered by pseudotime; T1 is the first 50 cells (cluster 0, meristem/early anthers), and T37 is the latest stage (cluster 3, late anther).

### Gene regulatory trajectories of anther and pollen development

We used GENIE3 (Huynh-Thu *et al*., 2010) to exemplify the capacity of the snRNA-seq data to infer the dynamics of gene regulatory networks (GRNs) during plant development. We reconstructed the GRNs for all clusters that were identified as “anthers” and estimated the strength of interactions between known transcription factors (TFs) versus all expressed genes. For example, the Figure 2d shows the GRN for cluster 15 representing an early anther stage. In our analysis, one of the main master TF (with most interactions) was *ABORTED MICROSPORES* (*AMS*), an already known regulator of anther development. We investigated more in-depth the regulatory dynamics of this TF using our data, the predicted targets of *AMS* and the related TF genes *bHLH089, bHLH091* and *bHLH010* (28-29) were expressed in a highly dynamic manner (Fig. 2c,d). AMS target genes at early stages were functionally enriched in chromatin remodeling (e.g. *BRAHMA*; *SET DOMAIN PROTEIN 16*) and pollen development (*DIHYDROFLAVONOL 4-REDUCTASE-LIKE1; ATP-BINDING CASSETTE G26*) (Fig. 2e). Late targets included metabolic enzymes as well as genes associated with RNA-regulatory processes. Newly identified marker genes covered the full anther developmental trajectory and are candidates for further mechanistic analyses.

### Validation of cell type markers genes

To validate the clustering analysis and dynamic anther transcriptome trajectory, we assessed the expression patterns of genes using promoter::NLS-GFP reporter lines. We selected 10 previously uncharacterized genes predicted to be specific or preferentially expressed in one of the clusters (Fig. 3). Seven out of 10 selected genes showed a specific expression in line with predictions (Fig. 3, Supplementary Fig. 10). Specific expression in the floral meristem was observed for genes AT1G63100 and AT3G51740 from cluster 11 (Fig. 3b,c). Moreover, gene AT4G11290 from cluster 14 showed highly specific expression in the stigma (Fig. 3e). The genes *AT5G20030, AT5G08250, AT1G23520* and *AT2G16750* were expressed in anthers and showed stage-specific expression as predicted by our analysis shown in Fig. 2 (Fig. 3f-i, Supplementary Fig. 10a,c,d). Gene *AT5G08250* from cluster 7, the first cluster of anther lineage, showed very strong expression in young anthers from flower 16 to flower 18 (nomenclature according to TraVaDB; Fig. 3f, Supplementary Fig. 10a); *AT5G20030* from cluster 15, which is an ‘early anther’ cluster, showed a peak in expression in flower 12 (Fig. 3g, Supplementary Fig. 10b). *AT2G16750* from cluster 6, was expressed strongly in older anthers in flower 10 and flower 11 (Fig. 3h, Supplementary Fig. 10c). Finally, *AT1G23520* from cluster 3, the last cluster of anther lineage, was found to be expressed in old anthers in flower 6 to flower 8 (Fig. 3i, Supplementary Fig. 10d). On the other hand, *AT1G54500* was expressed in sepal primordia and developing sepals (Fig. 3d), indicating that it is not specific to meristems as predicted for cluster 5. *AT3G05570* and *AT2G38995* were found to be more broadly expressed (not shown).

**Figure 3:**
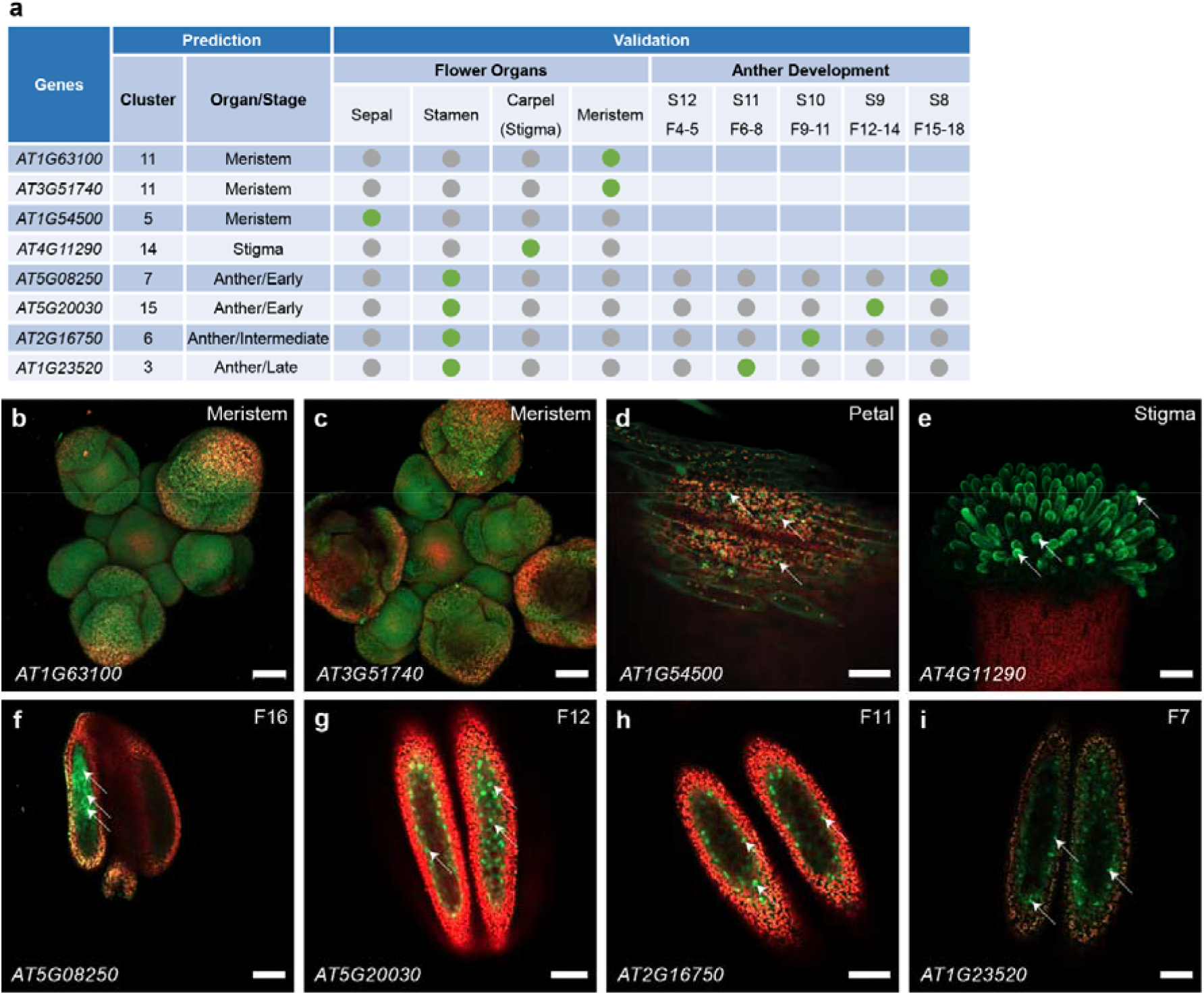
Validation of cluster-specific marker genes with transcriptional reporter lines. a) Summary of expression-specificity validation of selected marker genes. Green dots indicate positive and grey dots indicate negative GFP signals at particular developmental stages or flower organs. Flower numbers (F4-F18) are according to TraVaDB; flower developmental stages (S8-S12) are according to (Smyth *et al*., 1990). b) AT1G63100, meristem; c) AT3G51740, meristem; d) AT1G54500, sepal; e) AT4G11290, stigma; f) *AT5G08250*, anther from flower 16; g) *AT5G20030*, anther from flower 12; h) *AT2G16750*, anther from flower 11; i) *AT1G23520*, anther from flower 7. For the remaining confocal images see Supplementary Fig. 10. White arrowheads indicate exemplary GFP signals. Scale bars, 50 μm.

### Conclusions

Although protoplast isolation (PI) procedure may affect the plant transcriptome (Supplementary Fig. 1), it has been the main choice for plant single-cell sequencing and has been mostly applied to root samples so far (Zhang *et al*., 2019; Jean-Baptiste *et al*., 2019; Denyer *et al*., 2019; Shulse *et al*., 2019; Ryu *et al*., 2019). Here, we introduced a snRNA-seq methodology based in the efficient isolation of nuclei. Working with nuclei has the overall advantage of eliminating the dissociation-induced transcriptional responses, the compatibility with frozen samples and the possibility to carry out RNA sequencing from individual cells to study cell-types, like neurons, in which it is very difficult to recover intact cells (Grindberg *et al*., 2013; Bakken *et al*., 2018; Wu *et al*., 2019). More specifically for plants, the advantages in using nuclei include the elimination of the need to lysis the cell wall and the elimination of organelles and vacuoles. As a disadvantage, working with nuclei decreases the amount of RNA per individual cell and therefore potentially reducing the sensitivity of transcript detection.

Reporter gene analysis typically measures cytoplasmic RNA levels, while snRNA-seq measures nuclear expression, therefore potentially creating some difficulties validating predicted marker genes by reporter gene analysis. The nuclear isolation protocol is directly and easily applicable to a broad range of different plant tissues such as seedlings, flowers and leaves, and thus provides a versatile tool for plant single cell omics. In principle, various library preparation and sequencing methods can be combined with our generic nuclei isolation procedure.

Nanowell-based library preparation offered the possibility of visual quality control of individual nuclei, achieved high numbers of several thousand genes per cell and more than a thousand nuclei per run to sensitively detect plant cell (sub-) types. The number of nuclei can potentially be upscaled by using denser and/or larger nanowell-formats to further increase the number of nuclei for sequence analysis. The here applied nanowell-based approach resulting in deep cellular transcriptome data was of particular advantage to identify co-regulated genes and decipher gene networks underlying biological processes of interest. Along with the ever-growing range of nucleic acid sequencing technologies and plant genomics reference databases, single-nuclei genomics procedures are expected to become valuable tools to build maps of all plant cells of developing and adult tissues, and to measure cell-type-specific differences in environmental responses to gain novel mechanistic insights into plant growth and physiology (Rhee *et al*., 2019).

## EXPERIMENTAL PROCEDURES

### Preparation of plant tissues

One gram of *Arabidopsis thaliana* (Col-0) 10-day-old seedlings or 10 inflorescences were collected and snap-frozen in liquid nitrogen. The same procedure was applied for the following samples: 10 unopened buds of *Petunia hybrida* (W115), 8 unopened buds of *Antirrhinum majus*, 20 fully developed flowers and 1.3 g leaves of *Solanum lycopersicum* for testing the nuclei isolation pipeline.

### Preparation of nuclei

Frozen tissue was carefully crushed to small pieces in liquid nitrogen using a mortar and a pestle and transferred to a gentleMACS M tube that was filled with 5 ml of Honda buffer (2.5 % Ficoll 400, 5% Dextran T40, 0.4 M sucrose, 10 mM MgCl_2_, 1 µM DTT, 0.5% Triton X-100, 1 tablet/50 ml cOmplete Protease Inhibitor Cocktail, 0.4 U/µl RiboLock, 25 mM Tris-HCl, pH 7.4). This buffer composition enables efficient lysis of cell membranes while keeping the nuclei membranes intact (Moreno-Romero *et al*., 2017). The M tubes were put onto a gentleMACS Dissociator and a specific program (Supplementary Table 1) was run at 4 °C to disrupt the tissue and to release nuclei. The resulting suspension was filtered through a 70 µm strainer and centrifuged at 1000 *g* for 6 min at 4 °C. The pellet was resuspended carefully in 500 µl Honda buffer, filtered through a 35 µm strainer and stained with 3x staining buffer (12 µM DAPI, 0.4 U/µl Ambion RNase Inhibitor, 0.2 U/µl SUPERaseIn RNase Inhibitor in PBS). Nuclei were sorted by gating on the DAPI peaks using a BD FACS Aria III (200,000 – 400,000 events) into a small volume of landing buffer (4% BSA in PBS, 2 U/µl Ambion RNase Inhibitor, 1 U/µl SUPERaseIn RNase Inhibitor). Sorted nuclei were additionally stained with NucBlue from the Invitrogen Ready Probes Cell Viability Imaging Kit (Blue/Red), then counted and checked for integrity in Neubauer counting chambers. Quality of RNA derived from sorted nuclei was analyzed by Agilent TapeStation using RNA ScreenTape or alternatively by Agilent’s Bioanalyser 2100 system.

### Preparation of single-nucleus libraries using SMARTer ICELL8 Single-Cell System

The NucBlue and DAPI co-stained single-nuclei suspension (60□cells/µl) was distributed to eight wells of a 384-well source plate (Takara) and then dispensed into a barcoded SMARTer ICELL8 3’ DE Chip (Takara) by an ICELL8 MultiSample NanoDispenser (MSND, Takara). Chips were sealed and centrifuged at 500 g for 5 min at 4 °C. Nanowells were imaged using the ICELL8 Imaging Station (Takara). After imaging, the chip was placed in a pre-cooled freezing chamber, and stored at −80□°C for at least 2 h. The CellSelect software was used to support the identification of nanowells that contained a single nucleus. One chip yielded on average between 800 - 1200 nanowells with single nuclei. These nanowells were selected for subsequent targeted deposition of 50□nl/nanowell RT-PCR reaction mix from the SMARTer ICELL8 3’ DE Reagent Kit (Takara) using the MSND. After RT and amplification in a Chip Cycler, barcoded cDNA products from nanowells were pooled by means of the SMARTer ICELL8 Collection Kit (Takara). cDNA was concentrated using the Zymo DNA Clean & Concentrator kit (Zymo Research) and purified with AMPure XP beads. Afterwards, cDNA was used to construct Nextera XT (Illumina) DNA libraries followed by AMPure XP bead purification. Qubit dsDNA HS Assay Kit, KAPA Library Quantification Kit for Illumina Platforms and Agilent High Sensitivity D1000 ScreenTape Assay were used for library quantification and quality assessment. Strand-specific RNA libraries for sequencing were prepared with TruSeq Cluster Kit v3 and sequenced on an Illumina HiSeq 4000 instrument (PE100 run).

### Preparation of bulk RNA-seq libraries

Five 10-days-old *Arabidopsis thaliana* seedlings were collected into 1.5 ml screw-cap tubes with 5 glass beads, precooled in liquid nitrogen. Samples were homogenized by adding one half of the TRI-Reagent (Sigma-Aldrich, 1 ml per 100 mg) to each sample following sample disruption by using the Precellys 24 Tissue Homogenizer (Bertin) instrument for 30 sec and 4000 rpm. After homogenization, total RNA was extracted by adding the second half of the TRI-Reagent and the protocol was proceeded according to the manufacturer. To remove any co-precipitated DNA, a DNase-I digest was performed by using 1U DNase-I (NEB) in a total volume of 100 µl. Total RNA was cleaned-up by LiCl-precipitation using 10 µl 8 M LiCl and 3 vol. 100% ethanol incubating at -20 °C overnight. Following a spin down at 4 °C, 17,900 *g* for 30 min and 2 washing steps with 70% ethanol. The RNA pellet was dried on ice for 1 h and resuspended in 40 µl DEPC-water incubating at 56 °C for 5 min. Quality of total RNA was analyzed by Agilent TapeStation using RNA ScreenTape (Agilent) or alternatively by Bioanalyser 2100 (Agilent) system. Concentration was measured by a Qubit RNA BR Assay Kit (Thermo Fisher Scientific). One µg of total-RNA was used for RNA library preparation with TruSeq® Stranded mRNA Library Prep (Illumina), following the protocol according to the manufacturer. Quality and fragment peak size were checked by TapeStation using D1000 ScreenTape (Agilent) or alternatively by Bioanalyser 2100 (Agilent) system. Concentration was measured by the Qubit dsDNA BR Assay Kit (Thermo Fisher Scientific). Three replicates, composed of 5 seedlings each, were used separately throughout the whole procedure. Strand-specific RNA libraries were prepared using TruSeq Stranded mRNA library preparation procedure and the three replicates were sequenced on an Illumina NextSeq 500 instrument (PE75 run).

### Data pre-processing

Raw sequencing files (bcl) were demultiplexed and fastq files were generated using Illumina bcl2fastq software (v2.20.0). The command-line version of ICELL8 mappa analysis pipeline (demuxer and analyzer v0.92) was used for the data pre-processing and read mapping. Mappa_demuxer assigned the reads to the cell barcodes present in a predefined list of barcode sequences. Read trimming, genome alignment (*Arabidopsis thaliana* reference genome: TAIR10), counting and summarization were performed by mappa_analyzer with the default parameters. A report containing the experimental overview and read statistics for each snRNA-seq library was created using hanta software from the ICELL8 mappa analysis pipeline. The gene matrix generated by mappa_analyzer was used as input for the downstream analysis using R package Seurat v3 (Butler *et al*., 2018; Stuart *et al*., 2019).

### Quality control and data analysis

The analysis started by removing reads with barcodes representing the negative and positive controls included in all Takara Bio’s NGS kits. For the seedling samples, Seurat was used to filter viable nuclei by i) removing genes detected in less than 3 nuclei, ii) nuclei with less than 200 genes, iii) nuclei with more than 5% of reads mapped to mitochondria and iv) nuclei with more than 5% mapped to chloroplasts. Seurat *SCTransform* normalization method was performed for each one of the seedling replicates separately. Data from 3 seedling replicates were integrated using *PrepSCTIntegration, FindIntegrationAnchors* and *IntegrateData* functions. After running the *RunPCA* (default parameters), we performed UMAP embedding using *runUMAP* with *dims*=1:20. Clustering analysis was performed using *FindNeighbors* (default parameters) and *FindClusters* function with *resolution*=0.5. Differentially expressed genes were found using *FindAllMarkers* function and “wilcox” test, *logfc*.*threshold = 0*.*25* and *min*.*pct*=0.25. The sub-clustering analysis of root was performed using the *subset* function and the seedling clusters containing root cells (clusters: 3, 4, 6, 7, 9, 11 and 12; Fig. 1b). *SCTransform* and *RunUMAP* with *dims*=1:15 and *resolution*=1.5 were re-run after sub-setting the data and subsequently *FindAllMarkers* to find the differentially expressed genes across the sub-clusters, with the “wilcox” test, *logfc*.*threshold = 0*.*25* and using the RNA assay (normalized counts).

For the flower snRNA-seq dataset (900 nuclei), only genes encoded in the nucleus were used. Nuclei with i) less than 10,000 reads, ii) less than 500 genes containing 10 reads or iii) at least one gene covering more than 10% of the reads of a particular nucleus were filtered out. In addition, genes with less than 10 reads in at least 15 nuclei were also removed. The filtering step resulted in a dataset containing 856 nuclei and 14,690 genes. Seurat *SCTransform* normalization was applied to the filtered data using all genes as *variable*.*features*, and with parameters: *method*=“nb”, and *min_cells*=5. We used the *JackStraw* function in Seurat to estimate the optimal number of PCAs to be used in the analysis. After calculating the first 12 PCAs with *RunPCA*, we performed UMAP embedding using *runUMAP* with parameters *n*.*neighbors*=10, *min*.*dist*=.1, *metric*=“correlation” and *umap*.*method*=“umap-learn”. Clustering was done with *FindNeighbors* (default parameters) and *FindClusters* function using the SLM algorithm, *resolution*=1.15 and *n*.*iter*=100. Markers genes were found with the function *FindAllMArkers*, using the “wilcox” test and *min*.*pct*=0.25.

### Annotation

Annotation of the seedling and flower clusters was performed by visualizing the expression of the top 20 marker genes of each identified cluster on tissue and stage specific transcriptomes of TraVaDB (Transcriptome Variation Analysis Database, http://travadb.org, Klepikova *et al*., 2016).

### Reproducibility and correlation

To assess the reproducibility of our method, we compared the pooled number of reads overlapping each gene of each seedling replicate against one another in log2 space. The same was done to verify the similarity between unfixed and fixed seedling datasets.

The correlation between bulk and snRNA-seq datasets was investigated by comparing the average number of reads overlapping each gene in the snRNA-seq against bulk RNA-seq datasets. snRNA-seq (unfixed) and bulk RNA-seq of seedlings were generated in 3 biological replicates. Expression of bulk RNA-seq data was quantified with RSEM (Li and Dewey, 2014).

### Network analysis

GENIE3 (Huynh-Thu *et al*., 2010) was used to infer gene networks starting from the normalized expression data obtained from Seurat for each cluster independently, using the parameters *nTrees*=1000, and using as regulators the list of DNA binding proteins obtained from TAIR (www.arabidopsis.org). Genes expressed in less than 33% of the nuclei in a particular cluster were removed. Only the top 10,000 interactions were kept. Gene regulators with less than 10 predicted targets were also removed. Dynamics of the gene network through anther development were obtained by the following approach: first, all nuclei were ordered by their estimated developmental pseudotime using Monocle 3 (Trapnell *et al*., 2014) and cluster 0 (meristem/Early anther) as root cluster. Second, gene networks were estimated with GENIE, as described previously, using groups of non-overlapping sets of 50 nuclei that were previously ordered by its developmental pseudotime.

### Generation and Confocal Imaging of Reporter Lines

To validate expression specificity of the marker genes from our snRNA-seq approach, promoter::NLS-GFP (nuclear localization signal-green fluorescent protein) reporter lines were generated. The marker genes for validation were chosen from the pool of cluster-specific marker genes (p<0.05) that were not previously characterized in the literature (unknown marker genes). The genomic promoter region upstream of the ATG and until the closest neighboring gene was amplified by PCR and introduced into the entry vector pCR8:GW:TOPO by TA cloning (primers used for PCR are listed in Supplementary Table 3. Afterwards, the LR reactions were performed with the binary vector pGREEN:GW:NLS-GFP (Smaczniak *et al*., 2017) to generate GFP transcriptional fusions to a nuclear localization signal (NLS) peptide. All reporter constructs were transformed into the Col-0 Arabidopsis background, and multiple independent lines per construct were analyzed under a Zeiss LSM800 laser-scanning confocal microscope. Different floral organs were dissected and screened for the GFP signal by confocal microscopy under 20× and 63× magnification objectives. Auto-fluorescence from chlorophyll was collected to give an outline of the flower organs. A 488-nm laser was used to excite GFP and chlorophyll and emissions were captured using PMTs set at 410–530 nm and 650–700nm. Z-stack screens were performed for the floral meristem and stigma tissues to give a 3D structure visualization.

## DATA STATEMENT

The snRNA-seq and bulk RNA-seq data have been deposited in the ArrayExpress database at EMBL-EBI under accession number E-MTAB-9174.

## ACKNOWLEDGEMENTS

We thank Solenne Bourdier and Dijun Chen for their support.

## SUPPORTING INFORMATION

### Supplementary Tables

**Suppl. Table 1**: Program steps for gentle tissue disruption and nuclei release on the gentleMACS Dissociator and the instrument-specific commands.

**Suppl. Table 2**: List of marker genes in *Arabidopsis thaliana* seedlings and flowers.

**Suppl. Table 3**: List of primers used for PCR.

### Supplementary Data

**Suppl. Data 1**: Summary and read statistics of the snRNA-seq and bulk RNA-seq data.

### Supplementary Figures

**Suppl. Figure 1**: Effect of protoplast isolation (PI) in a root scRNA-seq experiment (Denyer *et al*., 2019).

**Suppl. Fig. 2**: Nuclei isolation quality in different plant tissues/species.

**Suppl. Fig. 3**: Summary of snRNA-seq seedling datasets.

**Suppl. Fig. 4:** Reproducibility of snRNA-seq.

**Suppl. Fig. 5**: Annotation of seedling cell-clusters using TraVaDB dataset.

**Suppl. Fig. 6**: Single-nuclei transcriptome analysis of a subset of root cells derived from snRNA-seq seedling data.

**Suppl. Fig. 7**: Correlation between unfixed and fixed seedling samples.

**Suppl. Fig. 8:** Single-nuclei transcriptome analysis of *A*.*thaliana* flower development.

**Suppl. Fig. 9**: Temporal trajectory in the floral snRNA-seq dataset

**Suppl. Fig. 10**: Validation of cluster-specific marker genes with transcriptional reporter lines.

