## Supplementary material for "Single-nuclei RNA-sequencing of plant tissues": Suppl. Data 1

#### Summary of snRNA-seq

| Sample | Number of Nuclei | Number of Reads | Mean Reads/Nucleus | Mean Genes/Nucleus | Reads Mapped to Genome | Uniquely mapped reads (%) |
|---|---|---|---|---|---|---|
| Seedling-Repl1 | 1,269 | 313 M | 255 K | 3,174 | 90% | 68.52% |
| Seedling-Repl2 | 973 | 223 M | 240 K | 2,308 | 95% | 73.91% |
| Seedling-Repl3 | 1,106 | 205 M | 188 K | 2,926 | 96% | 72.71% |
| Seedling-Fixed | 1,209 | 245 M | 177 K | 2,270 | 86% | 82.11% |
| Flower | 996 | 250 M | 267 K | 3,309 | 90% | 78.18% |

#### Summary of Bulk RNA-seq

| Sample | Number of reads | Reads mapped to Genome | Uniquely mapped reads (%) |
|---|---|---|---|
| bulk-Rep1 | 61 M | 91.31% | 86.46% |
| bulk-Rep2 | 60 M | 96.17% | 90.94% |
| bulk-Rep3 | 53 M | 93.36% | 89.38% |

#### Detailed report of snRNA-seq

### Experimental Overview

| | |
|---|---|
| **Sample Types** | Neg_Ctrl (48) \| Pos_Ctrl (48) \| sample (1,173) |
| **Total Reads** | 313.67 M |
| **Barcoded Reads** | 299.85 M |
| **Fraction Barcoded Reads** | 0.96 |
| **Barcodes Identified** | 1,269 |
| **Reads per Barcode** | 236.29 K |

#### *Arabidopsis thaliana* - Seedlings Rep1

### Read Statistics

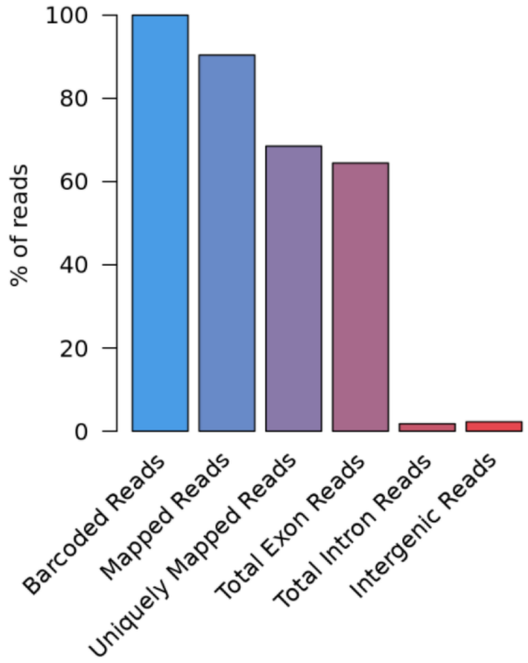


| | Read Count | % of Barcoded Reads |
|---|---|---|
| **Barcoded Reads** | 299,854,718 | 100.00 |
| **Trimmed Reads** | 289,379,784 | 96.51 |
| **Unmapped Reads** | 18,318,765 | 6.11 |
| **Mapped Reads** | 271,061,019 | 90.40 |
| **Uniquely Mapped Reads** | 205,452,448 | 68.52 |
| **Multimapped Reads** | 65,608,571 | 21.88 |
| **Total Exon Reads** | 193,263,409 | 64.45 |
| **Unique Exon Reads** | 182,430,105 | 60.84 |
| **Ambiguous Exon Reads** | 10,833,304 | 3.61 |
| **Total Intron Reads** | 5,375,695 | 1.79 |
| **Unique Intron Reads** | 5,322,651 | 1.78 |
| **Ambiguous Intron Reads** | 53,044 | 0.02 |
| **Intergenic Reads** | 6,813,344 | 2.27 |
| **Additional Information** | | |
| **Mitochondrial Reads** | 2,782,453 | 0.93 |
| **Ribosomal Reads** | 2,945,187 | 0.98 |

### Reads by Sample Type

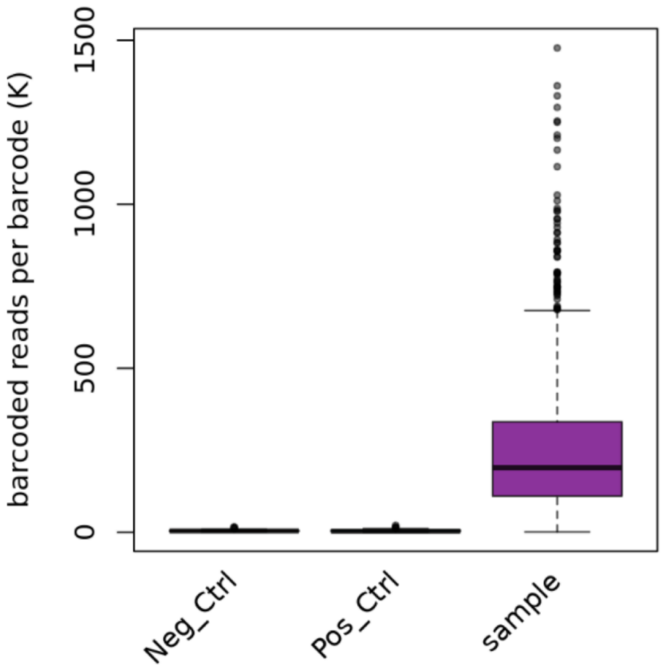


| | Reads per Barcode (Mean) | Reads per Barcode (Median) |
|---|---|---|
| **Neg_Ctrl** | 4,698.96 | 3,675.50 |
| **Pos_Ctrl** | 4,943.50 | 2,940.00 |
| **sample** | 255,236.04 | 196,529.00 |

### Genes by Sample Type

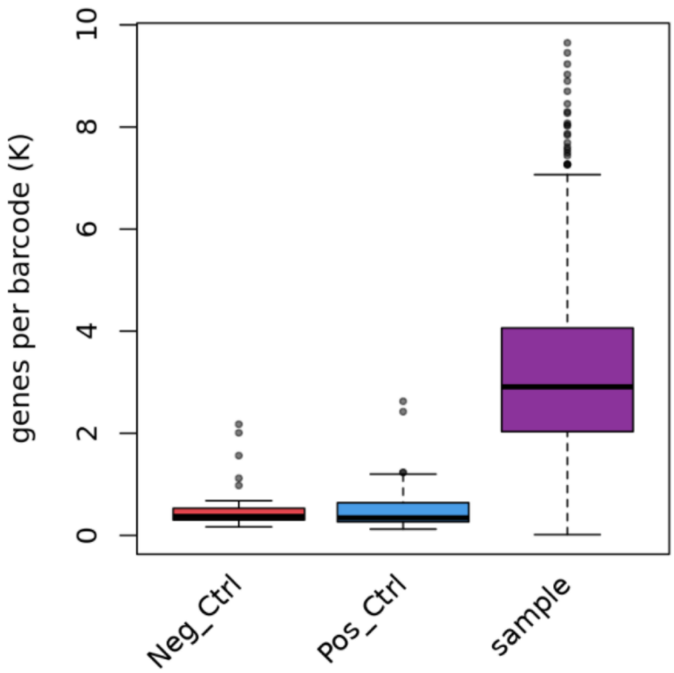


| | Genes per Barcode (Mean) | Genes per Barcode (Median) |
|---|---|---|
| **Neg_Ctrl** | 500.88 | 369.50 |
| **RPK 10000 > 0.1** | 500.85 | 369.50 |
| **RPK 10000 > 1.0** | 445.62 | 369.50 |
| **Pos_Ctrl** | 530.85 | 343.00 |
| **RPK 10000 > 0.1** | 530.79 | 343.00 |
| **RPK 10000 > 1.0** | 471.06 | 343.00 |
| **sample** | 3,174.58 | 2,909.00 |
| **RPK 10000 > 0.1** | 1,495.55 | 1,399.00 |
| **RPK 10000 > 1.0** | 664.22 | 632.00 |

### Experimental Overview

| | |
|---|---|
| **Sample Types** | Neg_Ctrl (47) \| Pos_Ctrl (47) \| sample (879) |
| **Total Reads** | 223.14 M |
| **Barcoded Reads** | 211.74 M |
| **Fraction Barcoded Reads** | 0.95 |
| **Barcodes Identified** | 973 |
| **Reads per Barcode** | 217.62 K |

### *Arabidopsis thaliana* - Seedlings Rep2

#### Read Statistics

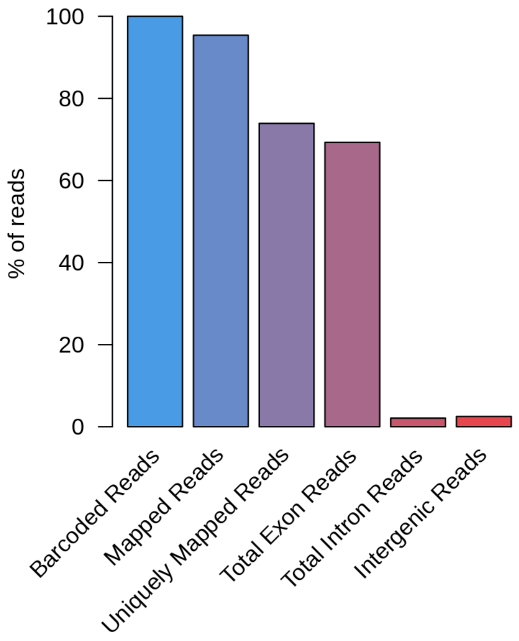


| | Read Count | % of Barcoded Reads |
|---|---|---|
| **Barcoded Reads** | 211,743,056 | 100.00 |
| **Trimmed Reads** | 207,173,647 | 97.84 |
| **Unmapped Reads** | 5,221,717 | 2.47 |
| **Mapped Reads** | 201,951,930 | 95.38 |
| **Uniquely Mapped Reads** | 156,500,615 | 73.91 |
| **Multimapped Reads** | 45,451,315 | 21.47 |
| **Total Exon Reads** | 146,729,979 | 69.30 |
| **Unique Exon Reads** | 139,064,104 | 65.68 |
| **Ambiguous Exon Reads** | 7,665,875 | 3.62 |
| **Total Intron Reads** | 4,432,812 | 2.09 |
| **Unique Intron Reads** | 4,396,736 | 2.08 |
| **Ambiguous Intron Reads** | 36,076 | 0.02 |
| **Intergenic Reads** | 5,337,824 | 2.52 |
| **Additional Information** | | |
| **Mitochondrial Reads** | 1,894,463 | 0.89 |
| **Ribosomal Reads** | 2,477,563 | 1.17 |

### Reads by Sample Type

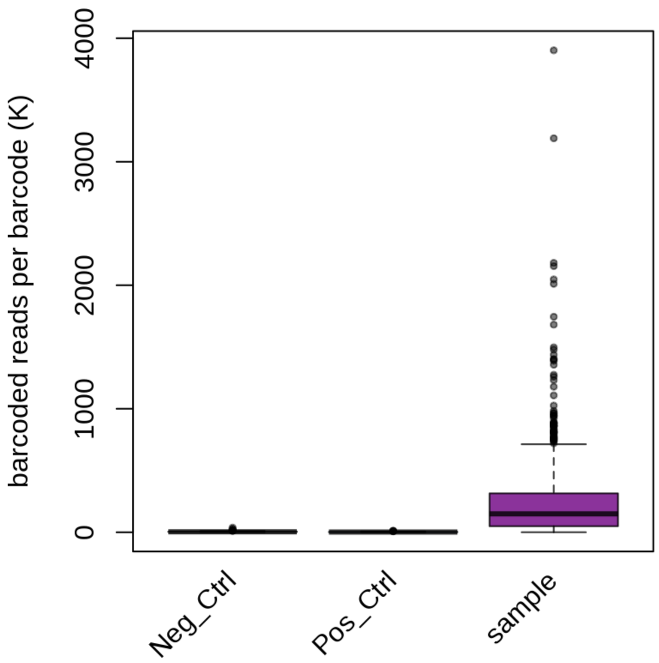


| | Reads per Barcode (Mean) | Reads per Barcode (Median) |
|---|---|---|
| **Neg_Ctrl** | 5,868.94 | 3,731.00 |
| **Pos_Ctrl** | 2,722.45 | 1,784.00 |
| **sample** | 240,431.47 | 149,274.00 |

### Genes by Sample Type

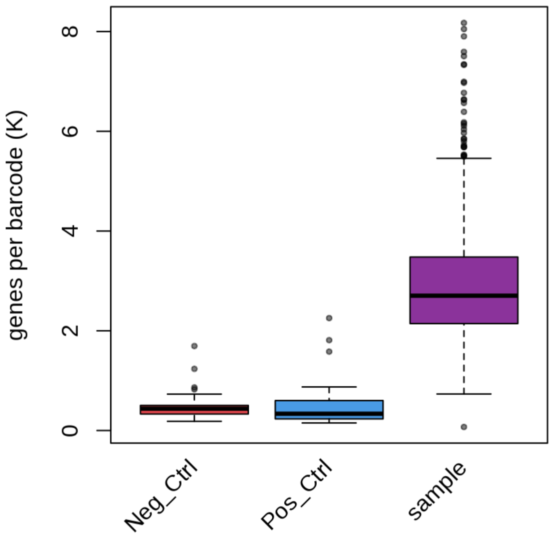


| | Genes per Barcode (Mean) | Genes per Barcode (Median) |
|---|---|---|
| **Neg_Ctrl** | 473.06 | 377.00 |
| **RPK 10000 > 0.1** | 473.06 | 377.00 |
| **RPK 10000 > 1.0** | 462.60 | 377.00 |
| **Pos_Ctrl** | 375.70 | 296.00 |
| **RPK 10000 > 0.1** | 375.70 | 296.00 |
| **RPK 10000 > 1.0** | 372.13 | 296.00 |
| **sample** | 2,308.36 | 2,147.00 |
| **RPK 10000 > 0.1** | 1,210.40 | 1,185.00 |
| **RPK 10000 > 1.0** | 596.03 | 567.00 |

### Experimental Overview

| | |
|---|---|
| **Sample Types** | Neg_Ctrl (48) \| Pos_Ctrl (47) \| sample (1,011) |
| **Total Reads** | 205.52 M |
| **Barcoded Reads** | 191.33 M |
| **Fraction Barcoded Reads** | 0.93 |
| **Barcodes Identified** | 1,106 |
| **Reads per Barcode** | 173.00 K |

*Arabidopsis thaliana* - Seedlings Rep3

### Read Statistics

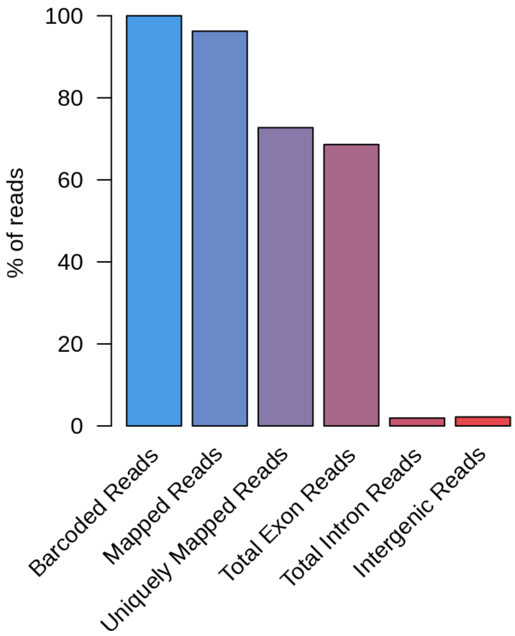


| | Read Count | % of Barcoded Reads |
|---|---|---|
| **Barcoded Reads** | 191,333,789 | 100.00 |
| **Trimmed Reads** | 186,940,462 | 97.70 |
| **Unmapped Reads** | 2,797,928 | 1.46 |
| **Mapped Reads** | 184,142,534 | 96.24 |
| **Uniquely Mapped Reads** | 139,124,356 | 72.71 |
| **Multimapped Reads** | 45,018,178 | 23.53 |
| **Total Exon Reads** | 131,251,219 | 68.60 |
| **Unique Exon Reads** | 124,316,795 | 64.97 |
| **Ambiguous Exon Reads** | 6,934,424 | 3.62 |
| **Total Intron Reads** | 3,688,398 | 1.93 |
| **Unique Intron Reads** | 3,642,567 | 1.90 |
| **Ambiguous Intron Reads** | 45,831 | 0.02 |
| **Intergenic Reads** | 4,184,739 | 2.19 |
| **Additional Information** | | |
| **Mitochondrial Reads** | 1,216,283 | 0.64 |
| **Ribosomal Reads** | 2,417,577 | 1.26 |

### Reads by Sample Type

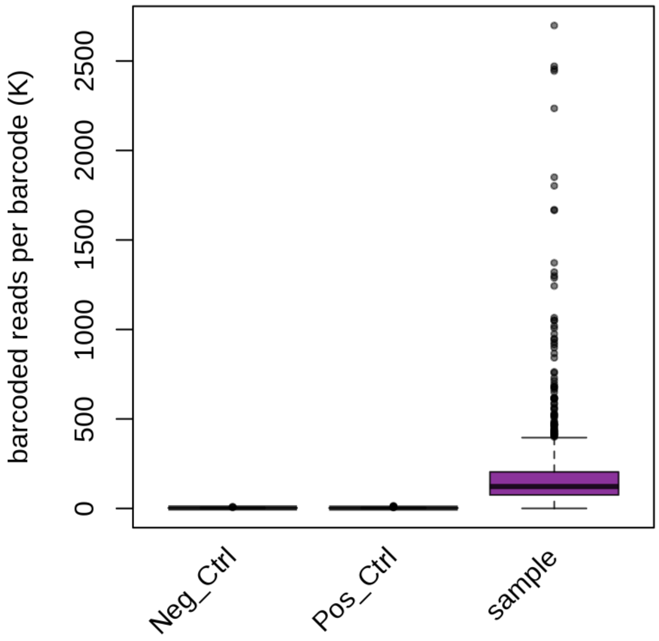


| | Reads per Barcode (Mean) | Reads per Barcode (Median) |
|---|---|---|
| **Neg_Ctrl** | 2,803.00 | 2,396.50 |
| **Pos_Ctrl** | 3,112.57 | 1,939.00 |
| **sample** | 188,974.24 | 122,326.00 |

### Genes by Sample Type

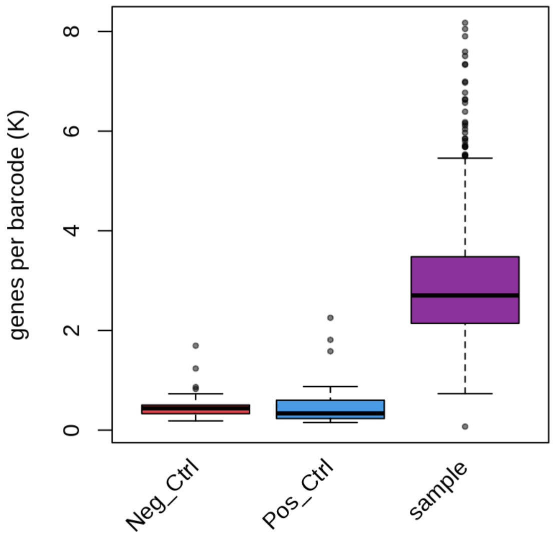


| | Genes per Barcode (Mean) | Genes per Barcode (Median) |
|---|---|---|
| **Neg_Ctrl** | 476.81 | 438.00 |
| **RPK 10000 > 0.1** | 476.81 | 438.00 |
| **RPK 10000 > 1.0** | 467.31 | 438.00 |
| **Pos_Ctrl** | 470.60 | 336.00 |
| **RPK 10000 > 0.1** | 470.60 | 336.00 |
| **RPK 10000 > 1.0** | 447.06 | 336.00 |
| **sample** | 2,926.01 | 2,703.00 |
| **RPK 10000 > 0.1** | 1,968.94 | 1,895.00 |
| **RPK 10000 > 1.0** | 879.87 | 855.00 |

### Experimental Overview

| | |
|---|---|
| **Sample Types** | Neg_Ctrl (45) \| Pos_Ctrl (45) \| sample (1,119) |
| **Total Reads** | 245.19 M |
| **Barcoded Reads** | 199.64 M |
| **Fraction Barcoded Reads** | 0.81 |
| **Barcodes Identified** | 1,209 |
| **Reads per Barcode** | 165.13 K |

#### *Arabidopsis thaliana* - Seedlings Fixed

### Read Statistics

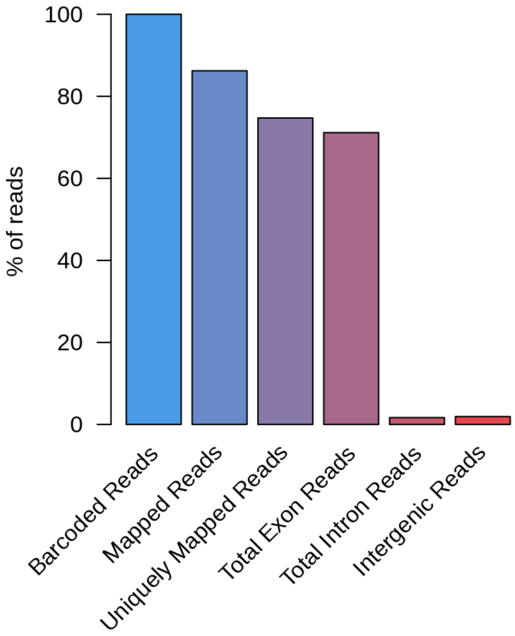


| | Read Count | % of Barcoded Reads |
|---|---|---|
| **Barcoded Reads** | 199,644,423 | 100.00 |
| **Trimmed Reads** | 181,669,195 | 91.00 |
| **Unmapped Reads** | 9,553,769 | 4.79 |
| **Mapped Reads** | 172,115,426 | 86.21 |
| **Uniquely Mapped Reads** | 149,160,816 | 74.71 |
| **Multimapped Reads** | 22,954,610 | 11.50 |
| **Total Exon Reads** | 142,028,753 | 71.14 |
| **Unique Exon Reads** | 134,859,300 | 67.55 |
| **Ambiguous Exon Reads** | 7,169,453 | 3.59 |
| **Total Intron Reads** | 3,321,079 | 1.66 |
| **Unique Intron Reads** | 3,301,467 | 1.65 |
| **Ambiguous Intron Reads** | 19,612 | 0.01 |
| **Intergenic Reads** | 3,810,984 | 1.91 |
| **Additional Information** | | |
| **Mitochondrial Reads** | 1,919,081 | 0.96 |
| **Ribosomal Reads** | 1,506,199 | 0.75 |

### Reads by Sample Type

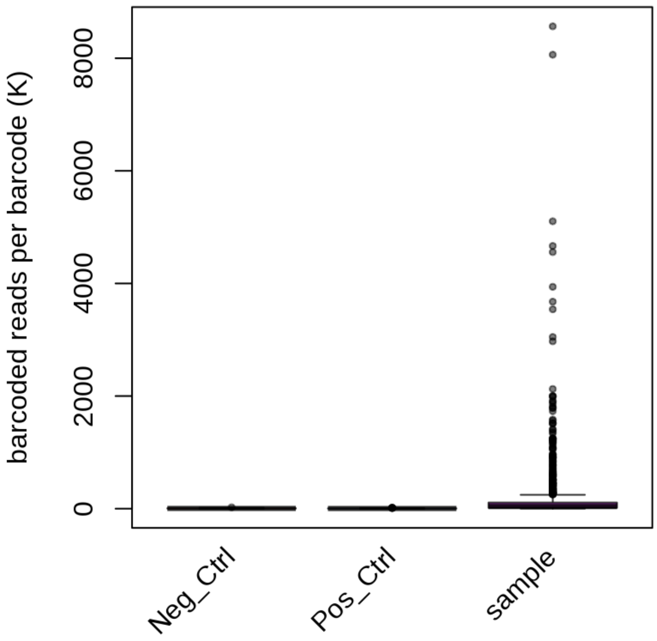


| | Reads per Barcode (Mean) | Reads per Barcode (Median) |
|---|---|---|
| **Neg_Ctrl** | 5,897.60 | 4,138.00 |
| **Pos_Ctrl** | 4,544.73 | 3,540.00 |
| **sample** | 177,993.31 | 37,876.00 |

### Genes by Sample Type

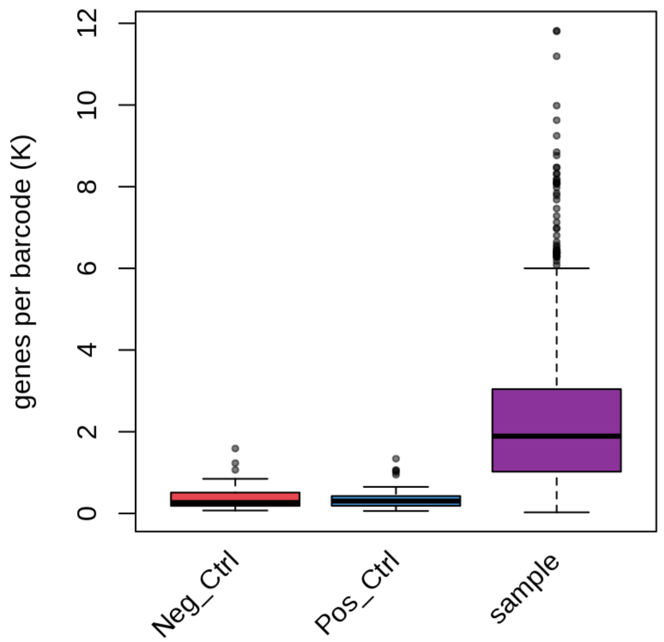


| | Genes per Barcode (Mean) | Genes per Barcode (Median) |
|---|---|---|
| **Neg_Ctrl** | 403.42 | 264.00 |
| **RPK 10000 > 0.1** | 403.42 | 264.00 |
| **RPK 10000 > 1.0** | 401.07 | 264.00 |
| **Pos_Ctrl** | 377.76 | 302.00 |
| **RPK 10000 > 0.1** | 377.76 | 302.00 |
| **RPK 10000 > 1.0** | 376.51 | 302.00 |
| **sample** | 2,270.27 | 1,892.00 |
| **RPK 10000 > 0.1** | 1,628.91 | 1,608.00 |
| **RPK 10000 > 1.0** | 794.85 | 796.00 |

### Experimental Overview

| | |
|---|---|
| **Sample Types** | Neg_Ctrl (48) \| Pos_Ctrl (48) \| sample (900) |
| **Total Reads** | 250.87 M |
| **Barcoded Reads** | 241.60 M |
| **Fraction Barcoded Reads** | 0.96 |
| **Barcodes Identified** | 996 |
| **Reads per Barcode** | 242.57 K |

#### *Arabidopsis thaliana* - Mixed flower stages

### Read Statistics

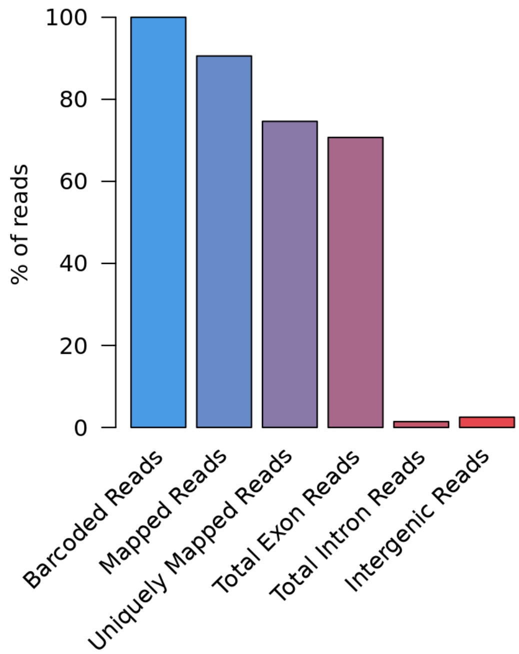


| | Read Count | % of Barcoded Reads |
|---|---|---|
| **Barcoded Reads** | 241,604,100 | 100.00 |
| **Trimmed Reads** | 230,595,698 | 95.44 |
| **Unmapped Reads** | 11,814,320 | 4.89 |
| **Mapped Reads** | 218,781,378 | 90.55 |
| **Uniquely Mapped Reads** | 180,275,535 | 74.62 |
| **Multimapped Reads** | 38,505,843 | 15.94 |
| **Total Exon Reads** | 170,801,636 | 70.69 |
| **Unique Exon Reads** | 160,471,912 | 66.42 |
| **Ambiguous Exon Reads** | 10,329,724 | 4.28 |
| **Total Intron Reads** | 3,416,113 | 1.41 |
| **Unique Intron Reads** | 3,396,512 | 1.41 |
| **Ambiguous Intron Reads** | 19,601 | 0.01 |
| **Intergenic Reads** | 6,057,786 | 2.51 |
| **Additional Information** | | |
| **Mitochondrial Reads** | 4,793,657 | 1.98 |
| **Ribosomal Reads** | 2,729,610 | 1.13 |

### Reads by Sample Type

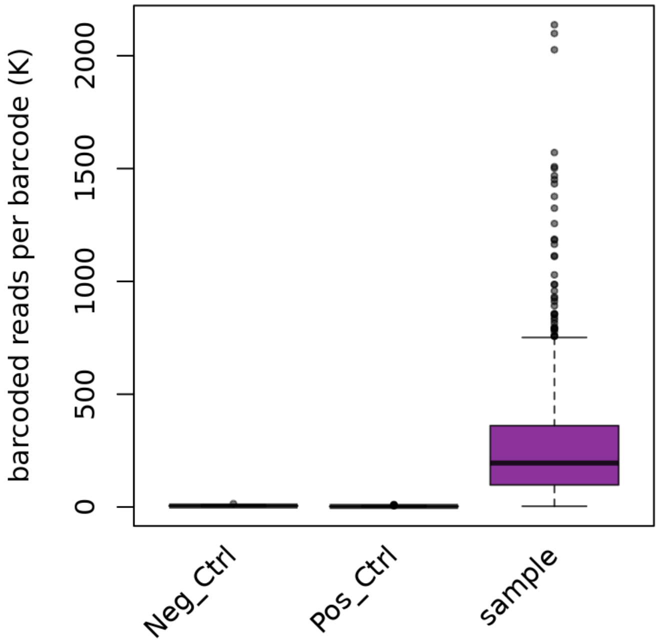


| | Reads per Barcode (Mean) | Reads per Barcode (Median) |
|---|---|---|
| **Neg_Ctrl** | 5,381.25 | 4,761.00 |
| **Pos_Ctrl** | 3,129.08 | 2,524.50 |
| **sample** | 267,995.12 | 195,046.00 |

### Genes by Sample Type

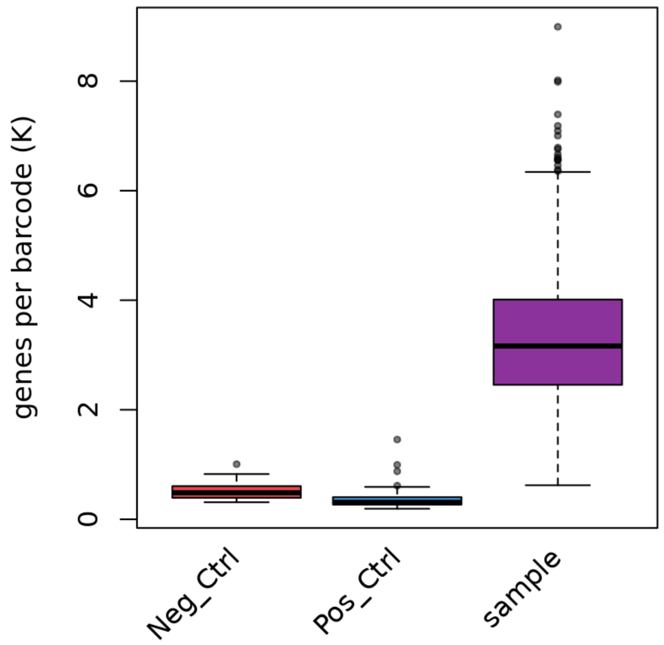


| | Genes per Barcode (Mean) | Genes per Barcode (Median) |
|---|---|---|
| **Neg_Ctrl** | 515.81 | 483.50 |
| **RPK 10000 > 0.1** | 515.81 | 483.50 |
| **RPK 10000 > 1.0** | 515.02 | 483.50 |
| **Pos_Ctrl** | 378.90 | 310.50 |
| **RPK 10000 > 0.1** | 378.90 | 310.50 |
| **RPK 10000 > 1.0** | 377.48 | 310.50 |
| **sample** | 3,309.21 | 3,164.50 |
| **RPK 10000 > 0.1** | 2,127.41 | 2,047.50 |
| **RPK 10000 > 1.0** | 1,143.84 | 1,126.00 |
