## Supplementary figures and images for "Single-nuclei RNA-sequencing of plant tissues"

### Suppl. Fig. 1

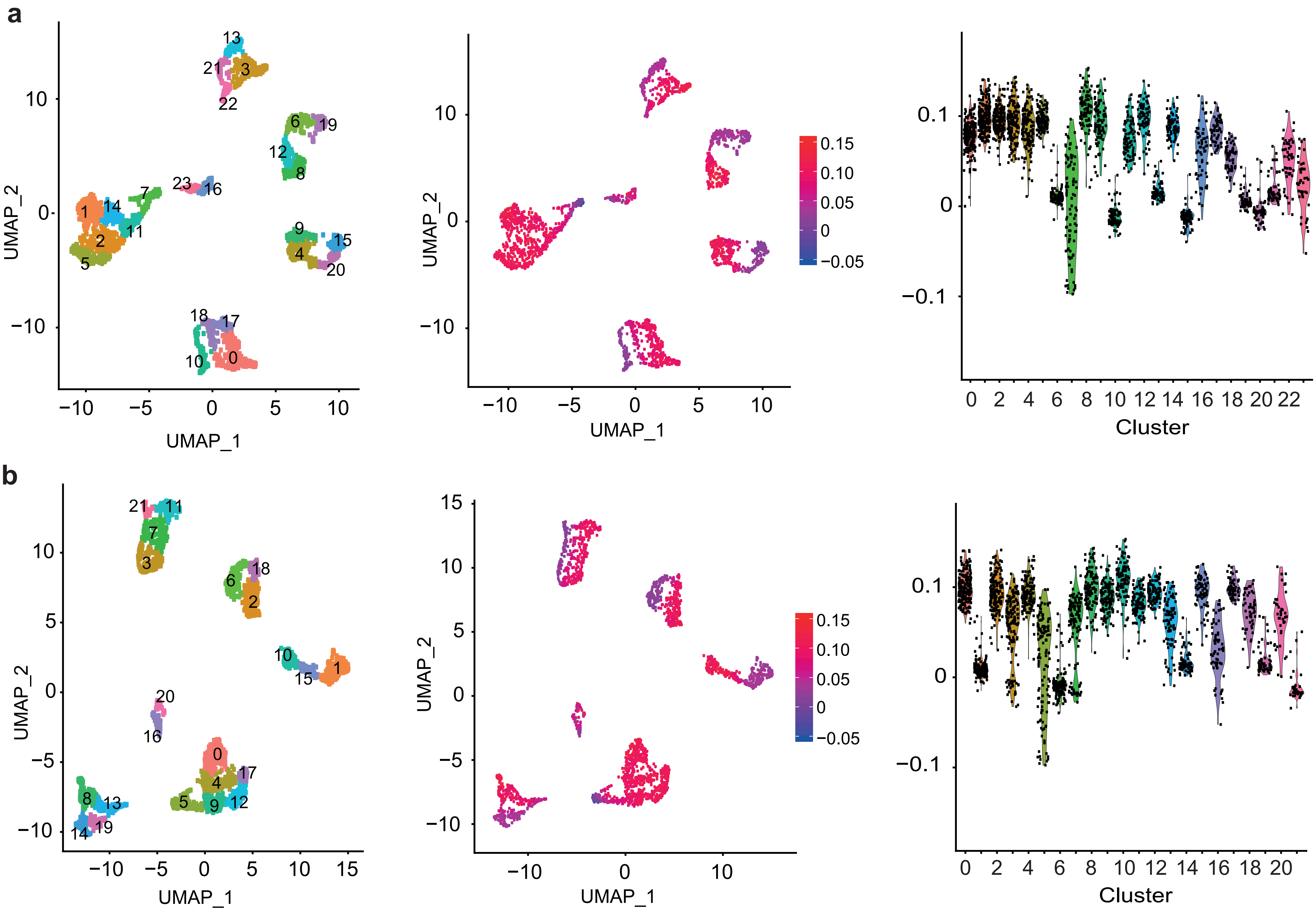

### Suppl. Fig. 2

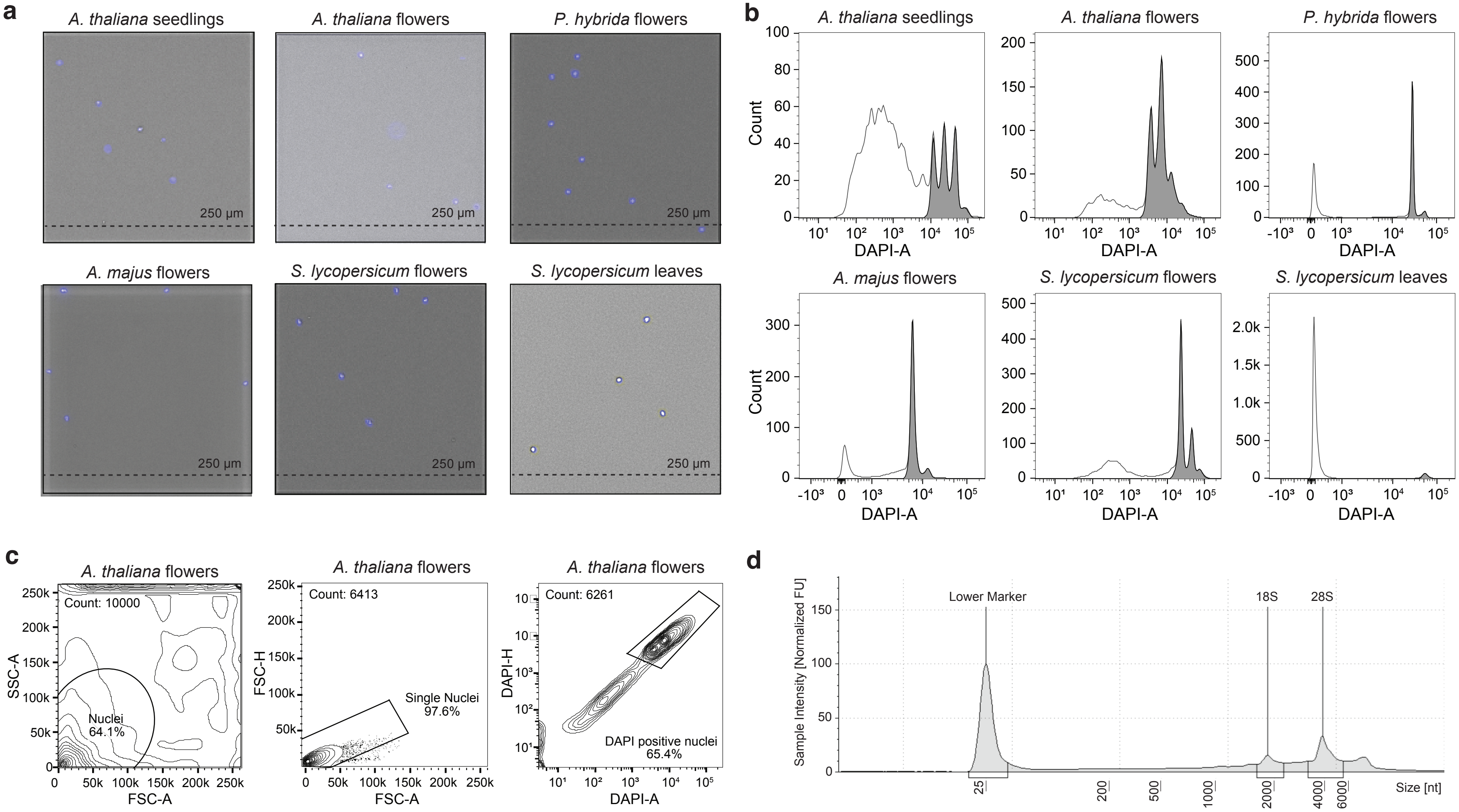

### Suppl. Fig. 3

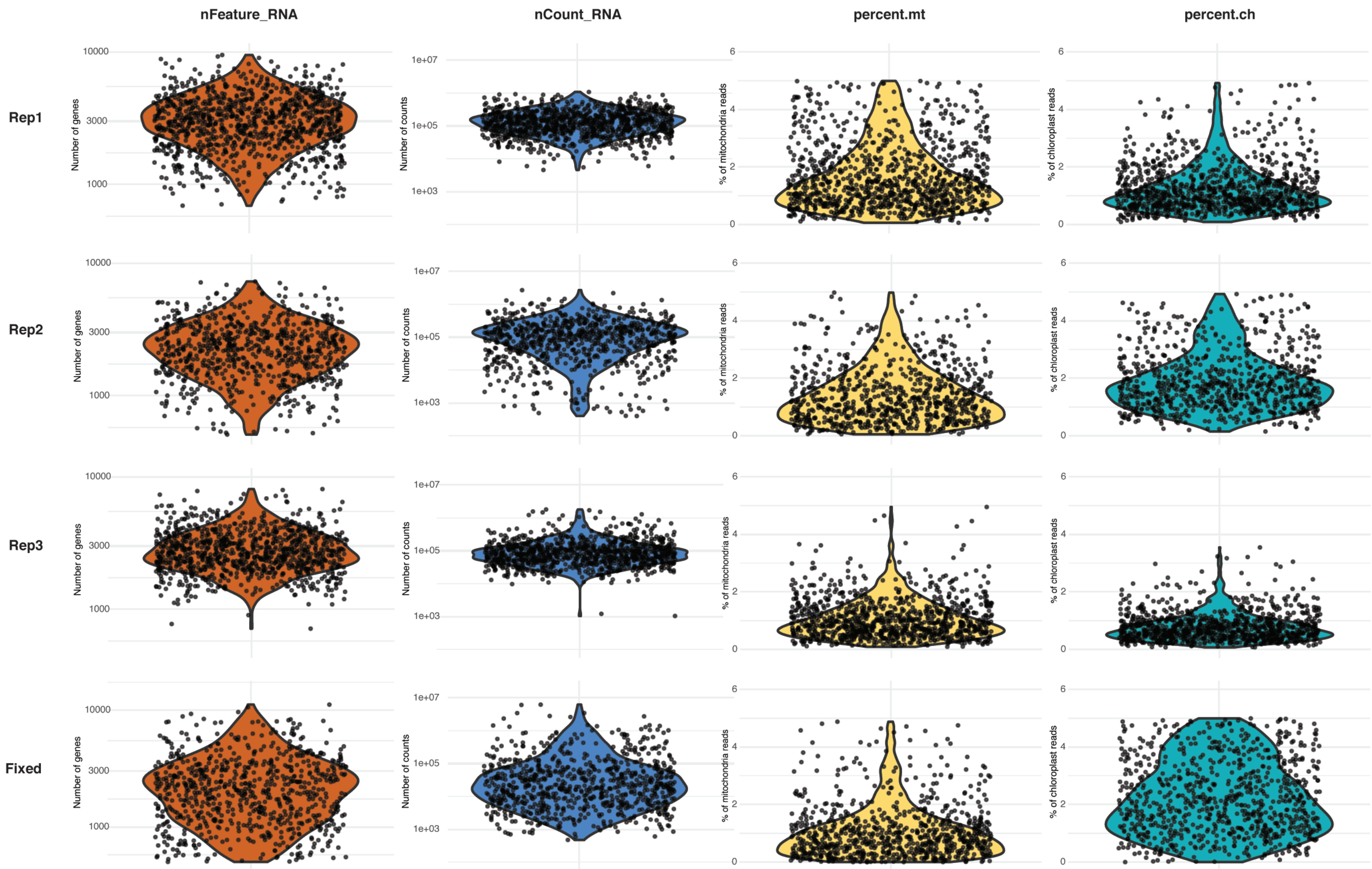

### Suppl. Fig. 4

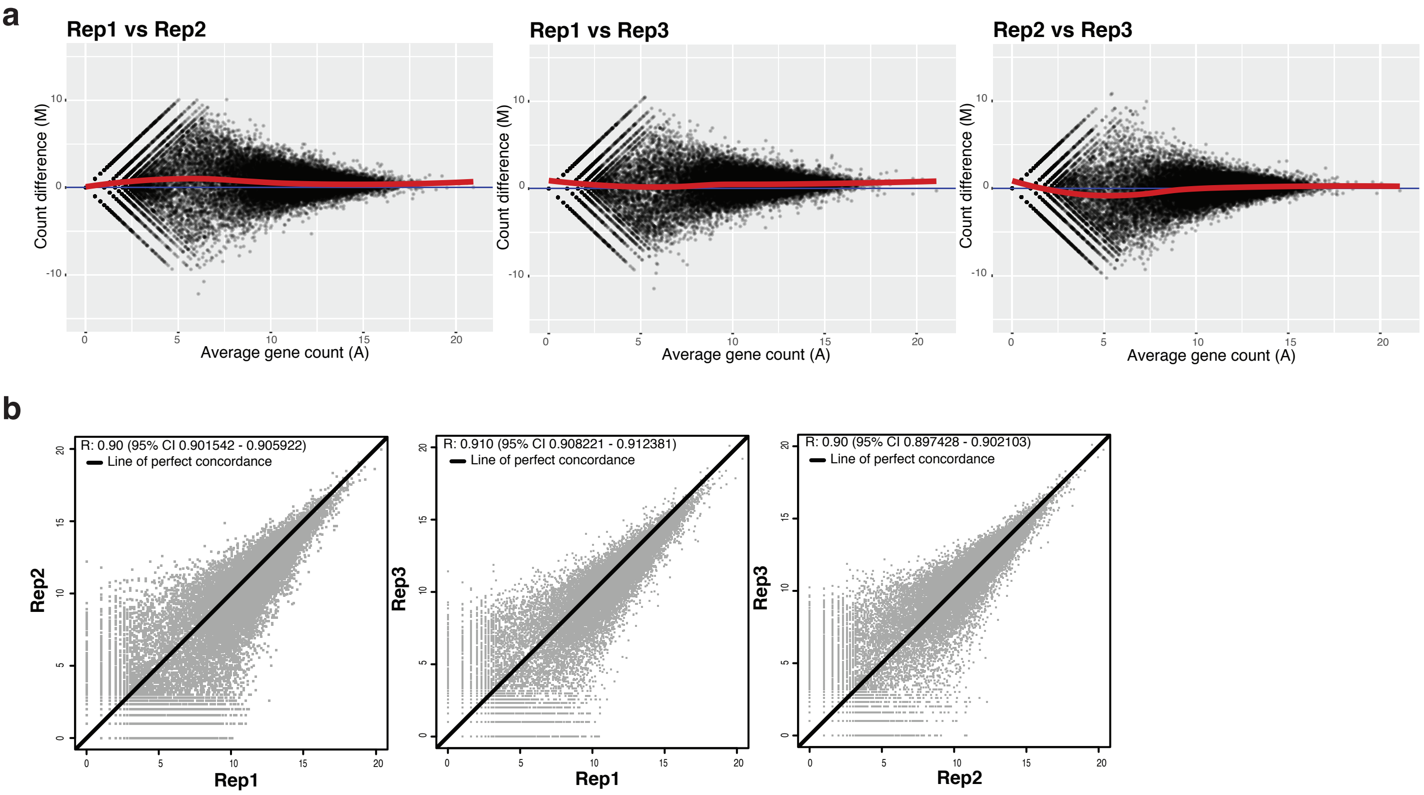

### Suppl. Fig. 5

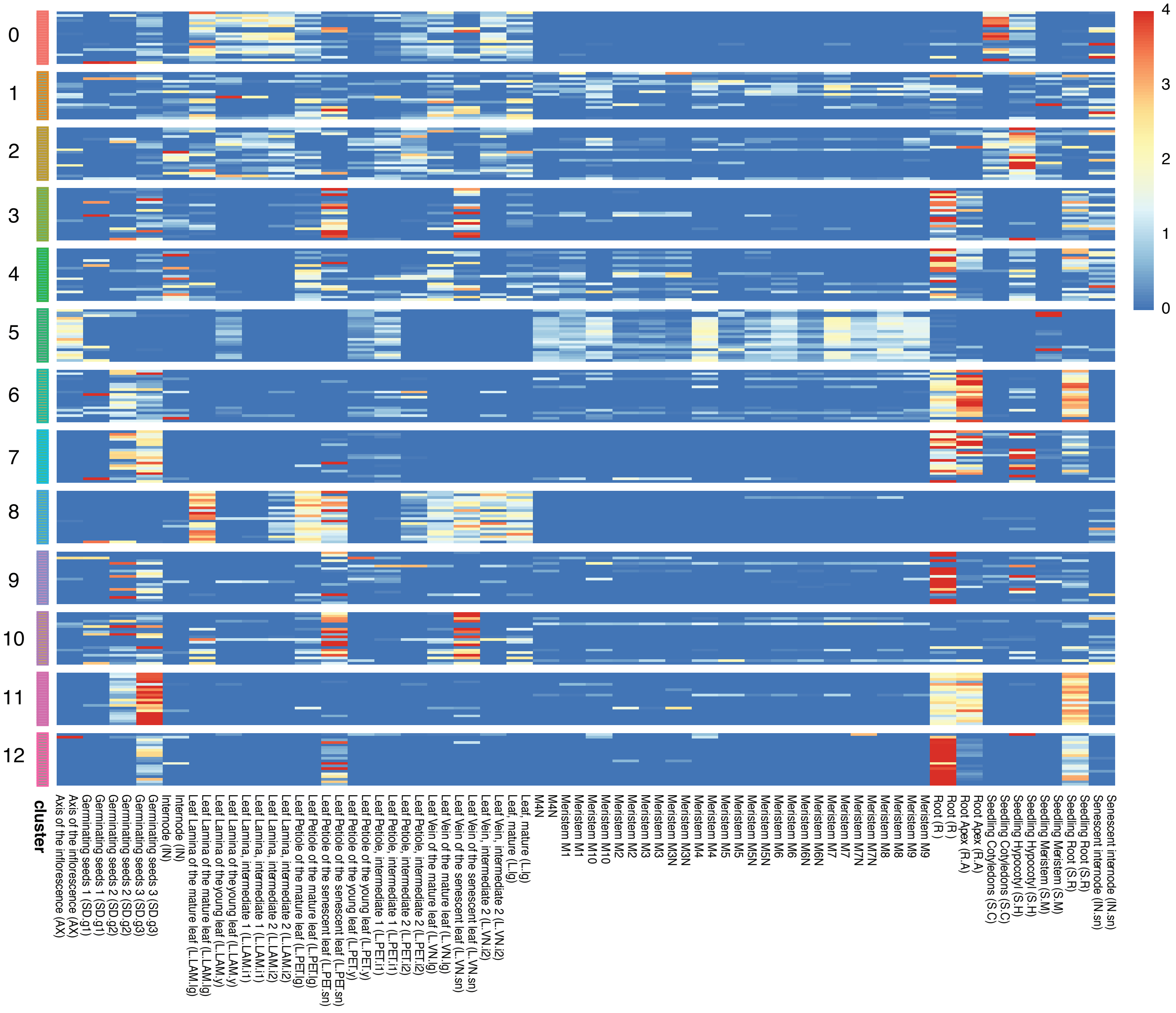

### Suppl. Fig. 6

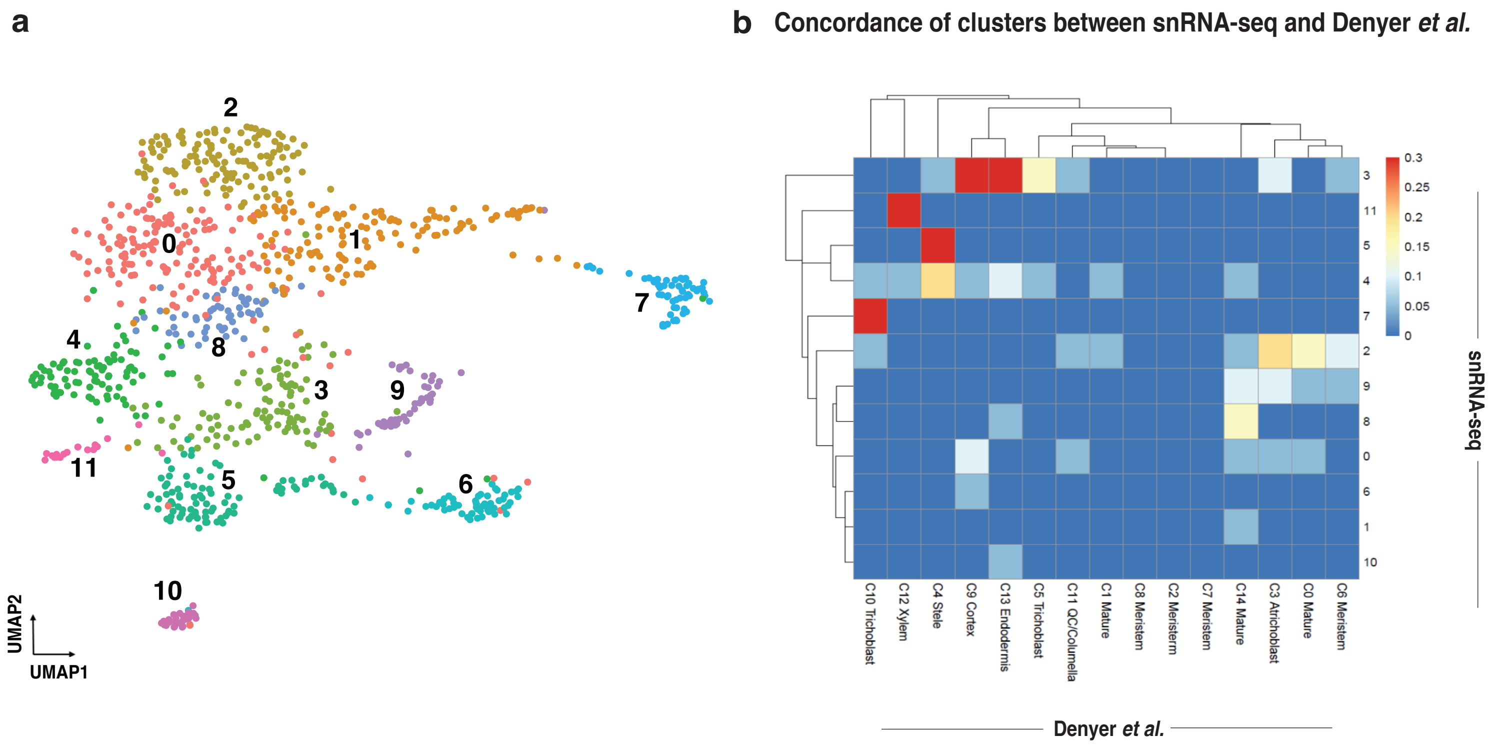

### Suppl. Fig. 7

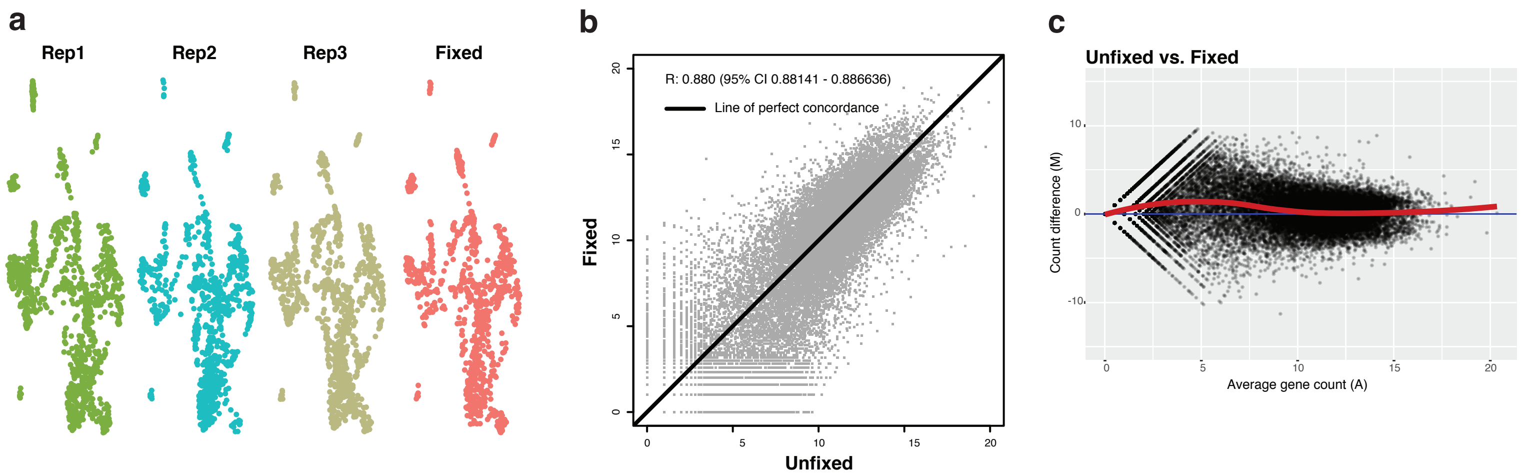

### Suppl. Fig. 8

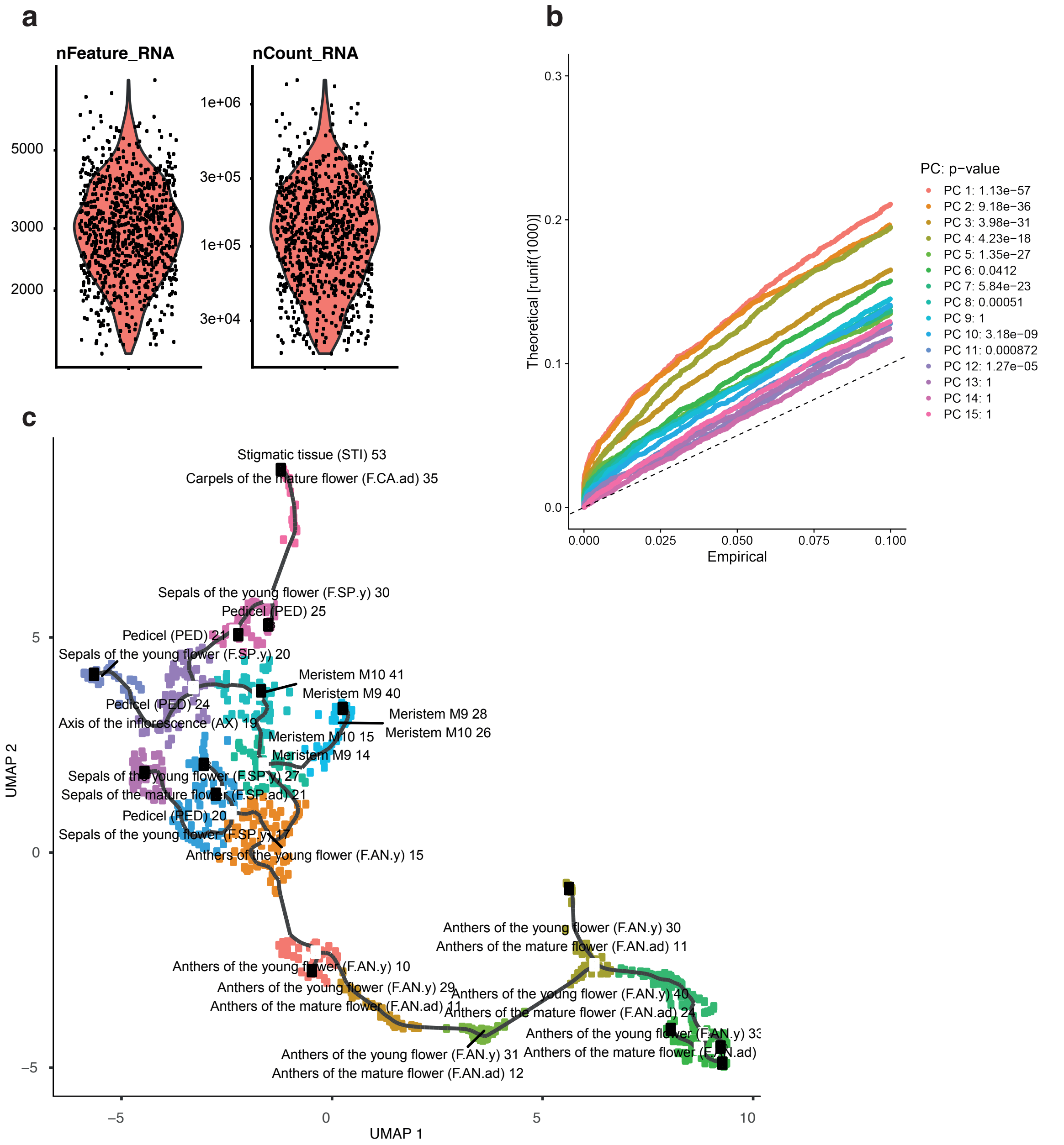

### Suppl. Fig. 9

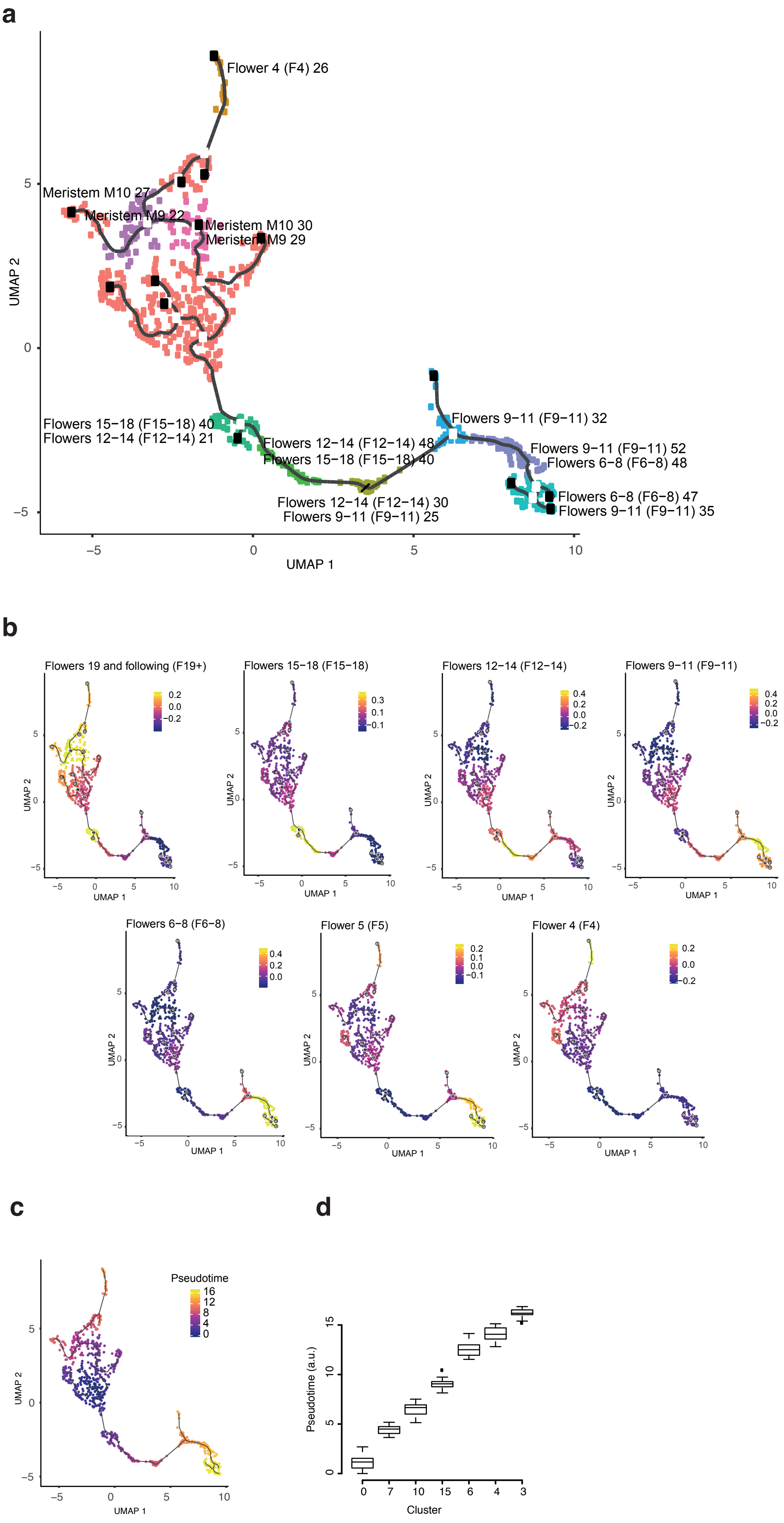

### Suppl. Fig. 10

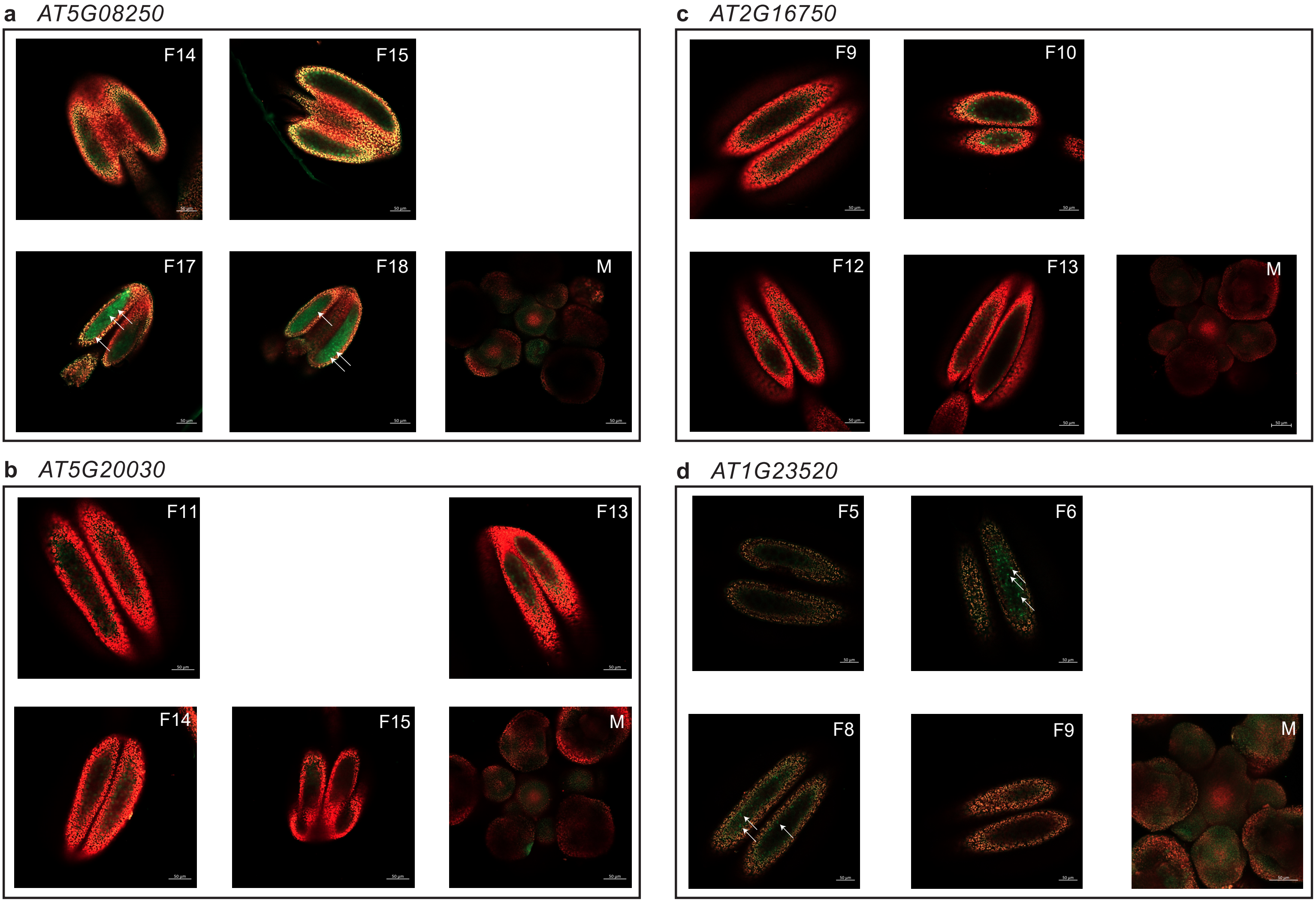
